# Domain organization and conformational change of dynactin p150

**DOI:** 10.1101/459040

**Authors:** Kei Saito, Takashi Murayama, Tomone Hata, Takuya Kobayashi, Keitaro Shibata, Saiko Kazuno, Tsutomu Fujimura, Takashi Sakurai, Yoko Y. Toyoshima

## Abstract

Dynactin is a principal regulator of the minus-end directed microtubule motor dynein. The sidearm of dynactin is essential for binding to microtubules and regulation of dynein activity. Although our understanding of the structure of the dynactin backbone (Arp1 rod) has greatly improved recently, structural details of the sidearm part remain elusive. Here, electron microscopy of individual molecules of the dynactin complex revealed that the sidearm was highly flexible and exhibited diverse morphologies. Utilizing mutants for nanogold labeling and deletion analysis, we determined the domain organization of the largest subunit p150 and identified a filamentous structure protruding from the head domain of the sidearm as the coiled-coil 1 (CC1), the dynein-binding domain, in p150. Furthermore, the protrusion formed by CC1 exhibited either a folded or an extended form, suggesting that CC1 works as an extending “arm”. These findings provide clues to understand how dynactin binds to microtubules and regulates dynein.

## Introduction

Dynactin is a multi-subunit complex that plays many essential roles in various cell functions, especially as an adaptor of dynein to vesicles or organelles (Kardon & Vale, 2009; Schroer, 2004). The importance of dynactin as a principal regulator of dynein and as an organizer of microtubule-based traffic is established but the molecular mechanisms of its diverse functions are not well known, mainly because of its very large and complicated architecture.

The dynactin complex is almost as large (~1.2 MDa) as cytoplasmic dynein and is composed of 11 different subunits (Schroer, 2004). This complex forms a unique asymmetric structure comprising two distinct domains, the Arp1 rod and the sidearm (Schroer, 2004). The Arp1 rod consists primarily of a polymer of Arp1 (Hodgkinson et al., 2005; Imai et al., 2006) and is responsible for cargo binding. One end of the rod (called the pointed-end) is capped by the “pointed-end complex” which consists of Arp11, p25, p27 and p62 (Eckley et al., 1999; Yeh et al., 2012; Yeh et al., 2013), whereas the other end (called the barbed-end) is capped by CapZ α/β (Schafer et al., 1994).

The sidearm is a thin elongated structure projecting from the barbed-end of the Arp1 rod. The main constituent of the sidearm is a dimer of p150*^Glued^* (hereafter called p150), which is the largest subunit in dynactin and interacts with dynein (Karki & Holzbaur, 1995; Vaughan & Vallee, 1995). The two globular heads are conspicuous at the distal sidearm and were thought to be formed by the N-termini of the p150 dimer, including CAP-Gly and the basic-rich domains (Schafer et al., 1994). The CAP-Gly domain binds to microtubules (MTs) (Waterman-Storer et al., 1995) and +TIPs (Steinmetz & Akhmanova, 2008), and particular mutations in this domain cause neuronal diseases (Farrer et al., 2009; Puls et al., 2003). The basic-rich domain directly alters the affinity of dynactin to MTs both *in vitro* and *in vivo* (Culver-Hanlon et al., 2006; Dixit et al., 2008; Kobayashi et al., 2017; Zhapparova et al., 2009). p150 is predicted to have a long (~330 aa) coiled-coil domain (CC1) almost immediately after the basic-rich domain and another coiled-coil (CC2; ~130 aa) on the C-terminal side.

The proximal end of the sidearm is called the “shoulder”. This domain is considered to be composed of some part of p150, four copies of p50 (also called dynamitin) and two copies of p24 (Eckley et al., 1999). The shoulder is essential for tethering the sidearm to the Arp1 rod because overexpression of p50 disrupts the interaction between the sidearm and the rod (Echeverri et al., 1996; Jacquot et al., 2010; Melkonian et al., 2007).

Recently, a cryo-electron microscopy (EM) study (Urnavicius et al., 2015) revealed the detailed structure of the dynactin complex. The rigid backbone of the complex including the Arp1 rod and the shoulder domain was especially well resolved. In contrast, the distal part of the sidearm, corresponding to p150, was only observed under conditions where p150 docked to the Arp1 rod. This configuration is markedly different from previously observed deep-etch rotary shadowing EM images of the dynactin complex (Schafer et al., 1994). In addition, p150 has scarcely been observed in other 2D or 3D averaged data of dynactin complex (Hodgkinson et al., 2005; Imai et al., 2006; Imai et al., 2014; Chowdhury et al., 2015). This elusiveness of p150 is likely derived from the flexible nature of the protein, which makes it difficult to properly understand the structure-function relationship of the dynactin complex, especially the mechanical basis for the dynamic interaction of dynactin with dynein and MTs.

CC1 is a particularly intriguing domain in p150 because, despite its well-known biochemical characteristic as the dynein-binding domain (King et al., 2003; Morgan et al., 2011), its structure and function remain controversial. CC1 was previously proposed to locate along the sidearm and the Arp1 rod (Schroer, 2004). A recent cryo-EM study assigned CC1 to the structure extending from the head domain (Urnavicius et al., 2015). The study suggested that CC1 folds back at the distal end of the sidearm, which might correspond to the second projection occasionally observed in the rotary shadowing EM (Schafer et al., 1994). However, the docked configuration in the cryo-EM left some important physiological questions unsolved. For example, what configuration does dynactin take in the cell and could CC1 adopt an extended conformation when dynactin binds with MTs or dynein, like postulated in Cianfrocco et al. (2015) or Carter et al. (2016)? Furthermore, the results of *in vitro* assays which examined the effect of CC1 on dynein motility are conflicting (Ayloo et al., 2014; Culver-Hanlon et al., 2006; Kardon et al., 2009; Kobayashi et al., 2017; Tripathy et al., 2014), implying there exists some regulatory mechanism within CC1. Thus, determining the location and conformation of CC1 in the dynactin complex, which is not averaged and not docked, is central in our pursuit of dynactin in action.

Herein, we report the folding pattern and domain organization of p150 in a human recombinant dynactin complex by combining negative stain EM and nanogold labeling. Our “non-averaged” observations of individual molecules revealed that the sidearm was a remarkably flexible structure and adopted various morphologies. We determined the localization of CC1 and CC2 in the sidearm, under the condition that both domains did not dock to the Arp1 rod. CC1 protruded from the head domain and exhibited two forms, a folded and an extended form, implying that CC1 undergoes a large conformational change. We propose a new model of the dynactin sidearm with the CC1 “arm” for dynein binding, and discuss the regulatory role of CC1 for MT binding.

## Results

### EM images of the dynactin complex revealed the flexibility of an “undocked” sidearm

We purified the human recombinant dynactin complex for EM observation. A streptavidin binding peptide (SBP) was fused to one of the subunits (Figure 1–S1) and all the other subunit components of the dynactin complex were co-purified by affinity chromatography (Figure 1–S2). We firstly used a complex including recombinant p62 (SBP-p62 in Figure 1–S1), which locates at the pointed end of the Arp1 rod (Kitai et al., 2011; Schafer et al., 1994), to obtain an intact form of the dynactin sidearm. The negative stain EM images revealed the sidearm complex projecting from one end of the Arp1 rod (Figure 1A; Figure 1–S3A, left). An overall configuration of the sidearm and the orientation of the projection were remarkably diverse, implying the flexible nature of the sidearm. Nevertheless, the sidearm exhibited common morphological features among molecules (Figure 1–S3A, middle), which enabled us to divide the structure of the sidearm into three domains: two globular heads, a thin neck and a shoulder projecting from the barbed end of the Arp1 rod (Figure 1A, left cartoon). Using these features as the reference points (Figure 1–S3A, middle and right), we described the geometry of the sidearm (Figure 1B), which suggested its preferable orientation and the range of motion.

**Figure 1.**
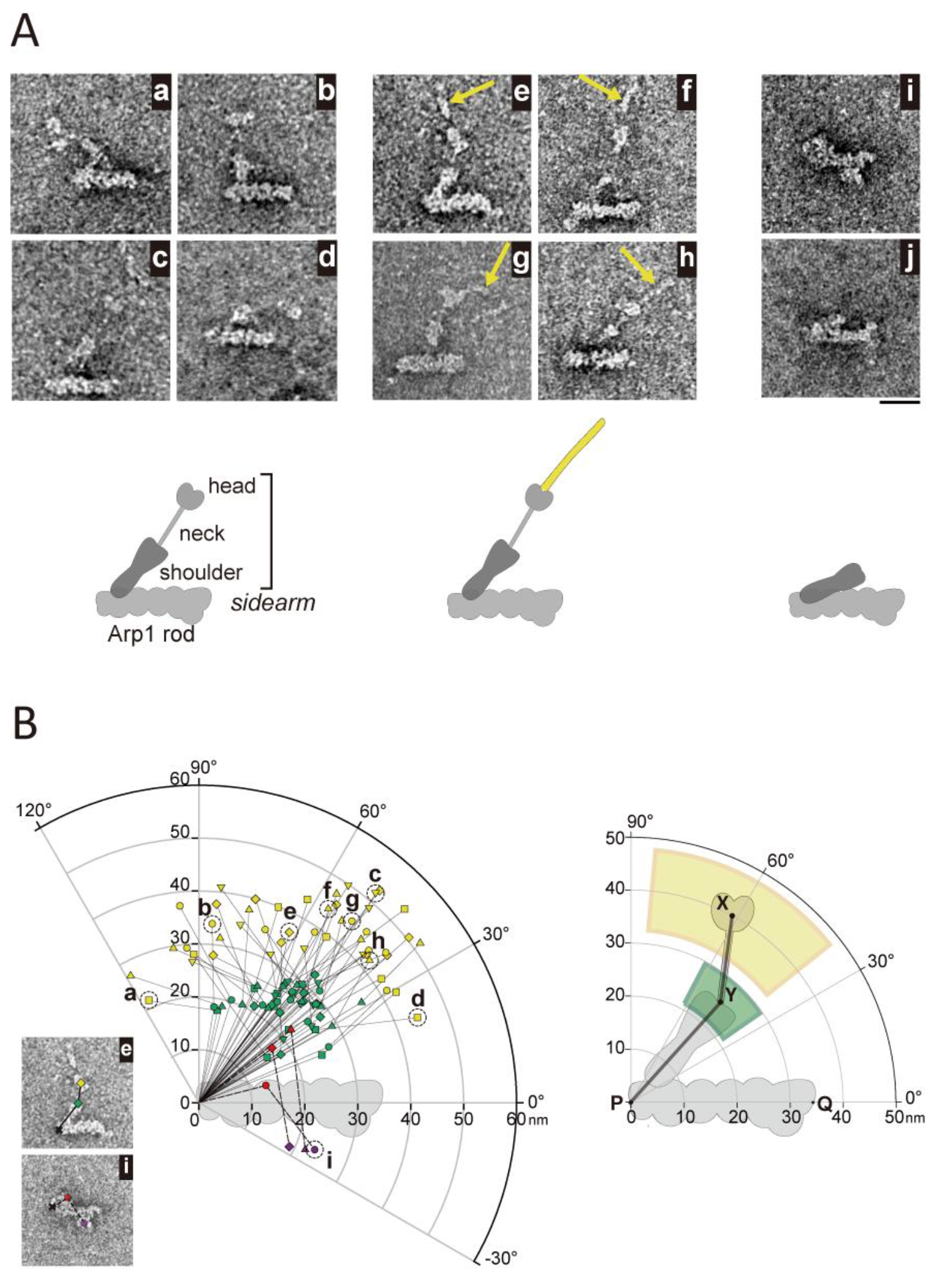
Negative stain EM of the human dynactin complex revealed morphological diversity of the sidearm and a thin filamentous structure at the tip. (A) Upper, negative stain EM images of the dynactin complexes (SBP-p62). The gallery shows the diverse morphologies of the sidearm, including the undocked sidearm (a)—(h) and the docked sidearm (i). The molecules in the panel (a)—(h) exhibits the distinguishable Arp1 rod, shoulder, neck and head domains; whereas the neck and the part of the shoulder is overlapped with the Arp1 rod in the panel (i). The head domain as well as the neck domain is not recognized in the panel (j). The panels (a), (b), (e) and (f) are inverted from the raw images to align the orientation of the Arp1 rod. Lower, cartoons to illustrate the morphological features and the configurations of the dynactin complex. (B) Quantitative comparison of the geometry of the sidearms. Left, the polar coordinates of the head and the neck-shoulder junction (base of neck) are shown. For the undocked sidearm, the yellow and the green plots indicate the positions of the head and base of the neck, respectively. For the docked sidearm, the purple and red plots indicate the positions of the head and base of the neck, respectively. For each molecule, the head and base of the neck are determined as illustrated in Figure 1—S3A, middle, and those positions are measured and mapped by setting the origin at the barbed end, to compare the orientation of the sidearm among molecules. See also Figure 1—S3A and Materials and methods for details. The data set is composed of dynactin molecules (SBP-p62) in which the reference points described above are clearly observed, with the undocked sidearm (yellow and green, *N*= 48) and the docked sidearm (purple and red, *N*= 3). Molecules in which the head domain was not recognized, such as the molecule (j) in (A), were not analyzed. Each polygonal line connecting the origin and the two plotted points indicates the backbone of the sidearm of each molecule (corresponding to the polygonal line X-Y-P in the right graph). To help distinguish between molecules, the sets of yellow and green (or red and purple) plots are marked with several symbols (circle, triangle up, triangle down, square, diamond). The plots labeled (a) to (i) correspond to the molecules shown in (A), and the EM images of (e) and (i) are shown here again as an example of the measurement. Right, the graph represents the mean and SD values of the head (yellow) and the base of neck (yellow) in the undocked form of the sidearm. The values are 40.0 ± 7.6 nm and 61.4 ± 23.1° for the head (X) (*N*= 48) and 25.4 ± 4.3 nm and 47.9 ±13.5° for the base of neck (Y) (*N*= 48). Bar represents 20 nm.

Morphological characteristics of the sidearm were similar to those of native chick dynactin revealed by deep-etch rotary shadowing EM (Schafer et al., 1994) in its overall configuration and flexibility as well as in the size and appearance of each domain, indicating that our recombinant approach retains the structural integrity of the dynactin complex. In our setting, the majority of the molecules exhibited an “undocked” form, in which the sidearm did not dock to the Arp1 rod (Figure 1A, **panels a-h**; Figure 1–S3A, left; Figure 1B, **yellow and green**) and a “docked” form of the sidearm reported by cryo-EM (Urnavicius et al., 2015) was rarely observed (Figure 1A, **panel i**; Figure 1B, **purple and red**). In a subset of molecules, the head and neck domains were not recognized as distinguishable structures (Figure 1A, panel j and right cartoon) and we excluded these molecules from the measurement in Figure 1B and subsequent analyses. We also found that glutaraldehyde fixation made the dynactin complex shrink. In this case, the undocked sidearm was rarely observed probably because of cross-linking (Figure 1–S3B). Therefore, we focused on the structures of the undocked sidearm without fixation hereafter, and examined the domain organization and conformation of the sidearm which retains its flexibility.

### Thin filamentous structure adjoining the head domain

Upon inspection of the head domain, we noticed that a thin filamentous structure adjoined the heads (Figure 1A, **panels e-h, yellow arrows**; Figure 1–S3C). This structure was particularly subtle compared with other parts of the dynactin complex and its visibility was rather varied even in the same grid; the ratio of the dynactin complex with the thin filamentous structure in the adequately stained region was 40-45% (*N* = 245, two independent experiments). The continuity between this structure and the two globular heads was not always distinct, possibly because of a particularly thin structure. Nonetheless, because it was found in almost every construct examined and mass spectrometry analysis of the purified sample detected no proteins known for binding with p150 (Table S1), we assumed that this filament was an essential part of the dynactin complex protruding from or associated with the head domain (Figure 1A, **middle cartoon**) but was not always visible by negative stain EM. The distance between the tip of the filament/protrusion and the center of the head was 28.8 ± 4.1 nm (mean ± SD, *N* = 20) and summarized with “somatometry” of the dynactin complex in Figure 1–S3D.

In previous studies, this structure was not observed in other negative stain EM observations (Imai et al., 2006; Imai et al., 2014; Chowdhury et al., 2015) but its appearance was similar to what was observed in the deep-etch rotary shadowed images (Schafer et al., 1994). Furthermore, it is probably identical to the structure assigned as CC1 in Urnavicius et al. (2015). Although their averaged image was produced in the docked form and unable to be directly compared with our images in the undocked form, the length was comparable between the two.

### Localization of the N- and C-termini of the p150 subunit

To dissect the molecular architecture of the dynactin sidearm, we made a series of His-tagged mutants of p150, p50 and p24 (Figure 1–S1) and labeled the tagged site in the dynactin complex with gold nanoparticles modified with Ni-NTA (see materials and methods). p150 is a large protein (~1250 aa) predicted to form two long coiled-coil structures (CC1 ~50 nm, CC2 ~20 nm). We further divided CC1 into CC1a and CC1b at the hinge point, where its coiled-coil structure is supposed to be disrupted (Figure 2A). p150 mutants carry the His-tag at respective locations: the N- and C-termini of p150, both sides and the hinge of CC1, and both sides of CC2 (Figure 2A, Figure 1–S1). The efficiency and specificity of nanogold labeling of the human dynactin used in this study has been reported previously (Kitai et al., 2011).

**Figure 2.**
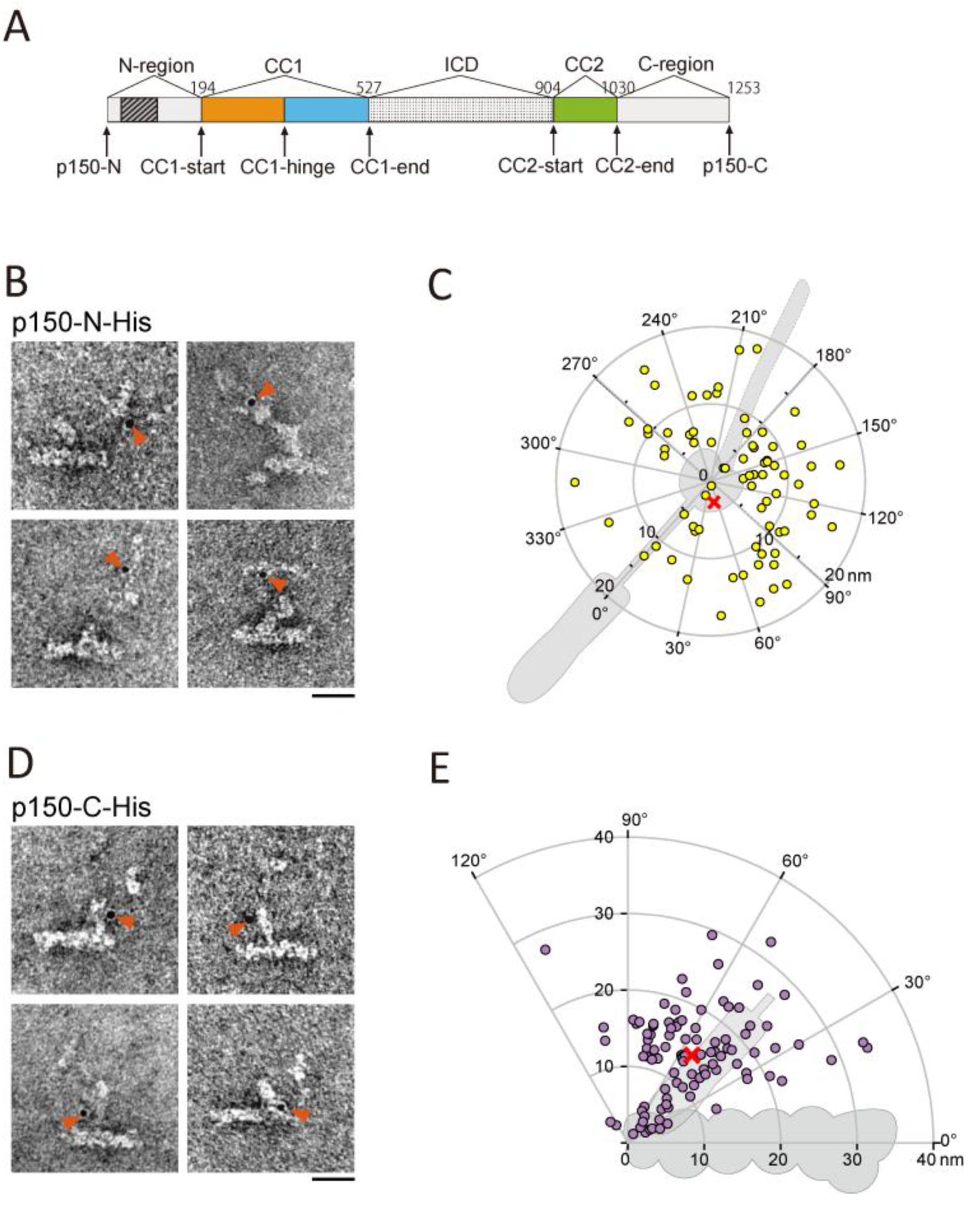
Localization of the N- and C-termini of p150 within the dynactin complex. (A) Diagram of the p150 sequence divided into five regions: N-region, CC1, ICD (inter coiled-coil domain), CC2 and C-region. The CAP-Gly domain (hatched), CC1a (orange), CC1b (light blue) and CC2 (light green) are indicated. The arrows indicate the sites where His-tags are inserted for nanogold labeling. (B) EM images of a mutant complex (p150-N-His) labeled with Ni-NTA gold nanoparticles (red arrowheads). (C) Distribution of the gold nanoparticles bound to p150-N-His. The polar coordinates of the centroid (red cross) are (2.8 nm, 51°) (*N*= 74). (D) EM images of a mutant complex (p150-C-His) labeled with the gold nanoparticles (red arrowheads). (E) The distribution of the gold nanoparticles bound to p150-C-His. The polar coordinates of the centroid (red cross) are (14.2 nm, 54°) (*N*= 98). Bars represent 20 nm.

The nanogold labeling of p150-N-His revealed that the N-terminus of p150 was localized around the head domain (Figure 2B; see also Table S2 for statistics of nanogold labeling experiments). Although the position of each gold nanoparticle was widely scattered, the centroid of the gold nanoparticles was located close to the center of the heads (Figure 2C). For p150-C-His, we found the gold nanoparticles around the shoulder (Figure 2D) and measured them from the barbed end of the Arp1 rod (Figure 2E). The location of the N-terminus is consistent with previous models (Schroer, 2004; Urnavicius et al., 2015). The location of the C-terminus does not agree with the model where the C-terminus of p150 extends along the Arp1 rod (Schroer, 2004) but agrees well with the interpretation of cryo-EM data that assigned one of the α-helices seen in the shoulder as p150 C-terminal structure (Urnavicius et al., 2015).

### The thin filamentous structure is formed by CC1 and the neck domain by CC2

Next, we focused on CC1, which is the longest coiled-coil in p150, a well-known dynein binding domain, and a supposed constituent of the filamentous structure adjoining the head domain (Urnavicius et al., 2015). The nanogold labeling experiments showed that CC1-start and CC1-end were located on the head (Figure 3A, **top and bottom**). The gold nanoparticles were found close to the junctions of the head and the filament (yellow arrows) in these two constructs. In contrast, CC1-hinge was located at the tip of the filament (Figure 3A, **middle**). Mapping of nanogold labeling results also showed that the plots for CC1-hinge exhibited a broader distribution and were more distant from the head than those for the CC1-start and the CC1-end (Figure 3B, Figure 3–S1). These results strongly indicate that CC1 folds back at the hinge.

**Figure 3.**
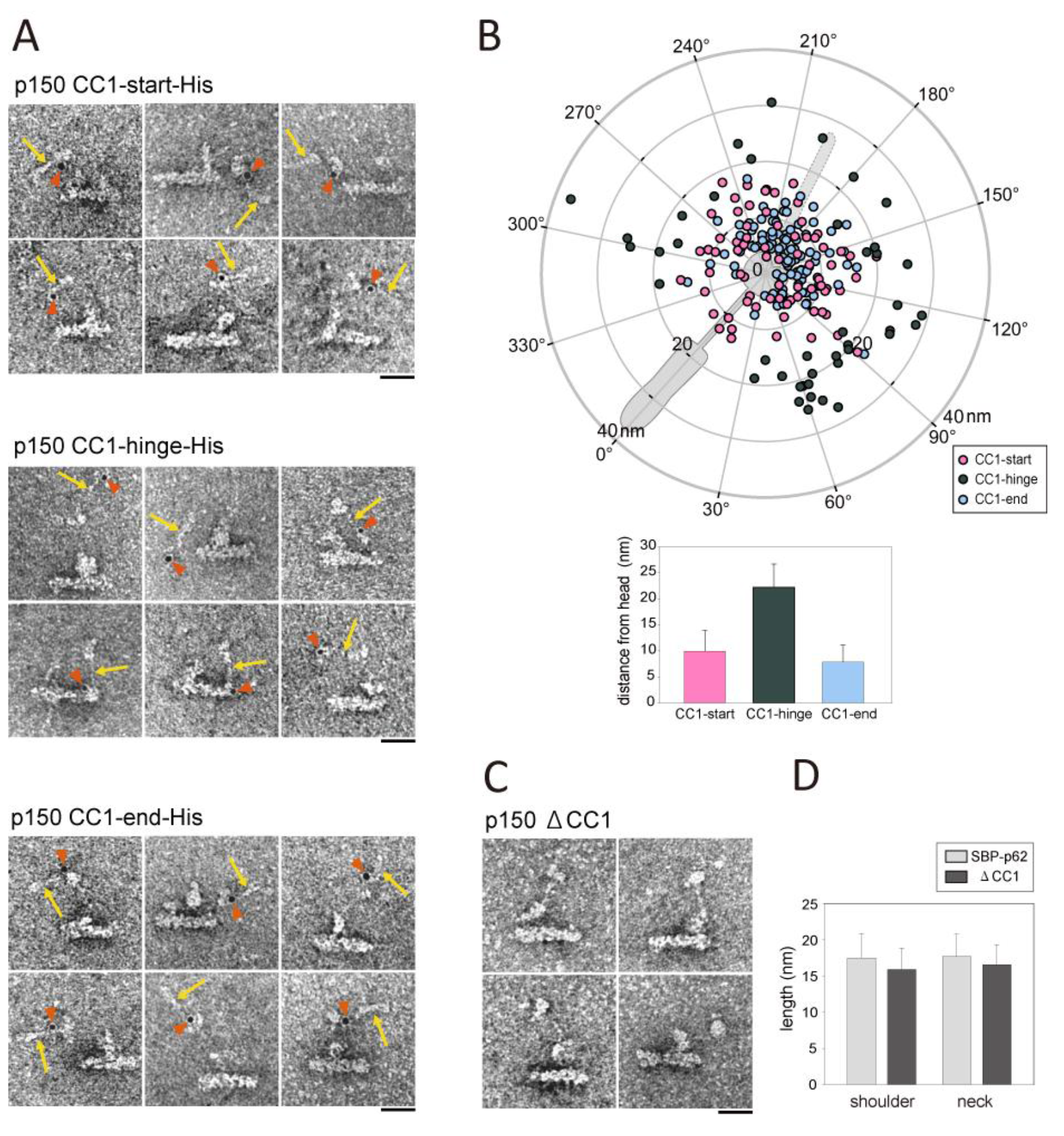
Identification of the p150 CC1 region. (A) EM images of the nanogold-labeled mutants, p150 CC1-start-His (top), p150 CC1-hinge-His (middle) and p150 CC1-end-His (bottom). Red arrowheads indicate the gold nanoparticles and yellow arrows indicate the thin filamentous structure. (B) Upper, the distribution of the gold nanoparticles bound to p150 CC1-start-His (pink) (*N*= 76), p150 CC1-hinge-His (dark green) (*N*= 41) and p150 CC1-end-His (light blue) (*N*= 110). See also Figure 3—S1 and Table S2 for the position of centroid and SD. Lower, the distance between the gold nanoparticles and the center of the head. Values are 10.0 ± 4.0 nm (*N*= 76), 22.2 ± 4.5 nm (*N*= 41) and 7.9 ± 3.3 nm (*N*= 110) (mean ± SD) for p150 CC1-start-His, p150 CC1-hinge-His and p150 CC1-end-His, respectively. (C) Negative stain EM images of the CC1 deleted p150 (p150 ΔCC1) dynactin complex. (D) Comparison of the lengths of the shoulder and the neck between p150 ΔCC1 and the wild type p150 (SBP-p62). The lengths of the shoulder (the segment YZ in Figure 1–S3D) of SBP-p62 and p150 ΔCC1 were 17.5 ± 3.3 nm (*N*= 50) and 15.9 ± 2.9 nm (*N*= 40), respectively (mean ± SD). The lengths of the neck (the segment XY in Figure 1–S3D) of SBP-p62 and p150 ACC1 were 17.7 ± 3.0 nm (*N*= 50) and 16.6 ± 2.8 nm (*N*= 40), respectively (mean ± SD) (Welch’s t-test, p = 0.02 for the shoulder and 0.06 for the neck). Bars represent 20 nm.

We subsequently made a CC1-deletion mutant of p150 (p150 ΔCC1 in Figure 1–S1). In the EM images, p150 ΔCC1 exhibited a similar configuration as SBP-p62 including wild-type p150 (Figure 3C, Figure 3–S2). The lengths of the shoulder and the neck were almost the same between ΔCC1 and wild-type p150 (Figure 3D). Collectively, we concluded that the thin filamentous structure was an essential component of the dynactin sidearm protruding from the head and it was formed by CC1 of p150 folding back at the hinge. Contrary to the previous model in which CC1 located along the sidearm and the Arp1 rod (Schroer, 2004), the recent cryo-EM study proposed that CC1 forms the structure projecting from the head (Urnavicius et al., 2015) and our results clearly support it.

The labeling of the second coiled-coil, CC2, showed that CC2-start was located at the head domain (Figure 4A,B) and CC2-end at the junction of the neck and shoulder domains (base of the neck) (Figure 4C,D). In addition, p150 ΔCC2 exhibited neck-lacking morphology, where the head and the shoulder were in close proximity, and the protrusion formed by CC1 was still evident (Figure 4E). These results demonstrated that CC2 was the neck itself, which also supports the model based on cryo-EM (Urnavicius et al., 2015).

**Figure 4.**
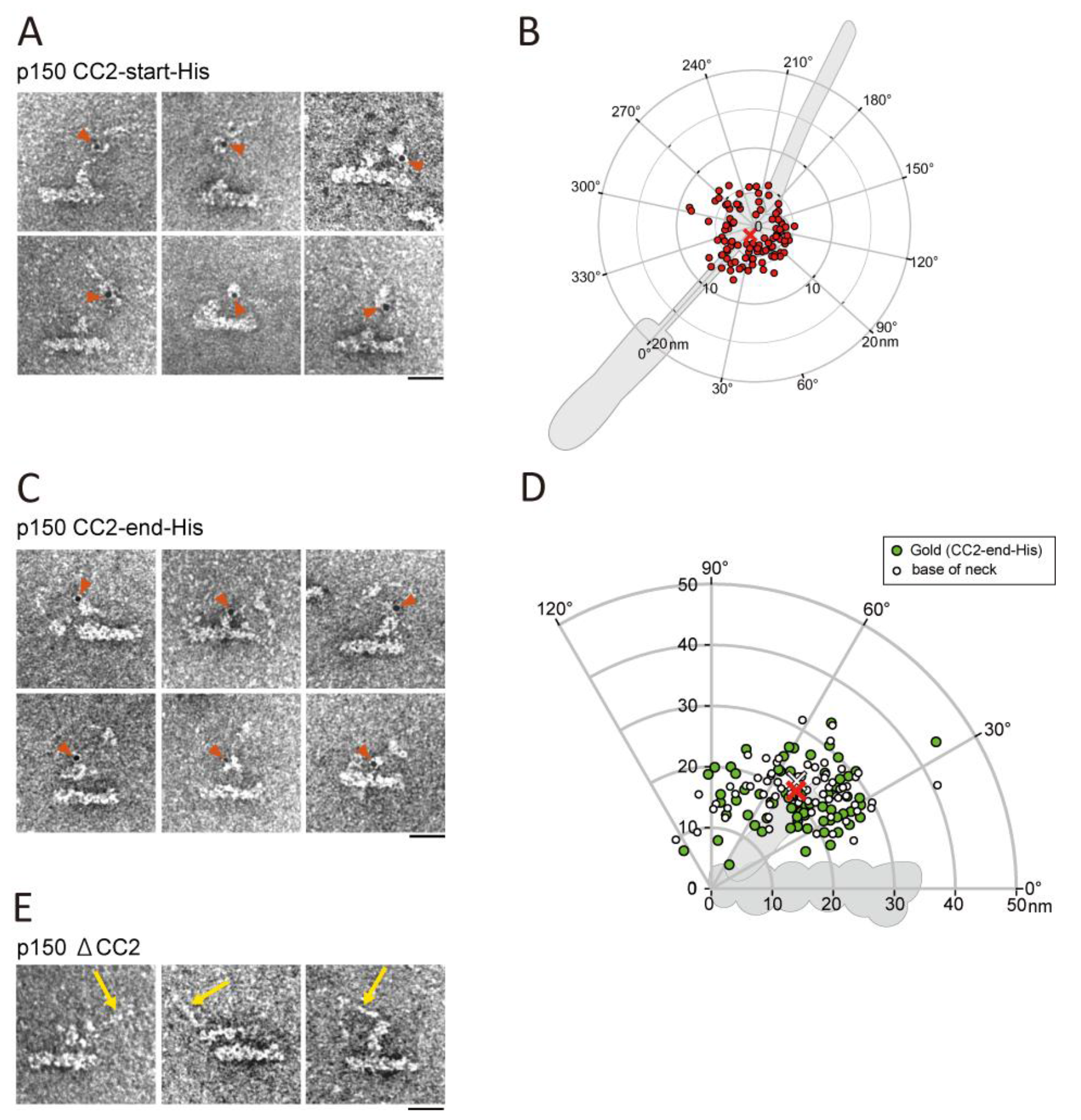
Identification of the p150 CC2 region. (A) EM images of p150 CC2-start-His labeled with the gold nanoparticles (red arrowheads). (B) The distribution of the gold nanoparticles bound to p150 CC2-start-His. The polar coordinates of the centroid (red cross) were (1.2 nm, 15°) (*N*= 95). (C) EM images of p150 CC2-end-His labeled with the gold nanoparticles (red arrowheads). (D) The distribution of the gold nanoparticles in the p150 CC2-end-His mutant (green circles) (*N*= 73). For comparison, the position of base of the neck (point Y in Figure 1—S3A) is shown for the same data set (white triangles). The red cross indicates the centroid of the gold nanoparticles (20.3 nm, 46°) and the white cross indicates the centroid of the base of neck (21.5 nm, 48°). (E) EM images of p150 ΔCC2. Yellow arrows indicate the protrusion formed by CC1. Bars represent 20 nm.

### C-region of p150 is indispensable for the complex assembly

Our nanogold labeling demonstrated that p150 C-terminus located at the middle of the shoulder (Figure 2D) and CC2-end at base of the neck (Figure 4D), which suggests that C-region of p150 (Figure 2A) resides in a part of the shoulder. The shoulder is thought to consist of p150, p50, and p24 subunits (Eckley et al., 1999). To clarify the molecular arrangement of the shoulder domain, we next focused on p50, p24 and C-region of p150.

The His-tag at the N-termini of p50 (p50-N-His) and p24 (p24-N-His) (Figure 1–S1) were labeled and found to be localized close to the barbed end of the Arp1 rod (Figure 5A, upper). The gold nanoparticles of the both mutants exhibited similar distributions (Figure 5B) and they were closer to the barbed end than p150-C-His (Figure 2D, Table S2). The mutants with the His-tag at the C-terminus (p50-C-His and p24-C-His) were purified as complexes (Figure 1–S2C). However, these constructs did not produce clear results. For p50-C-His, the gold nanoparticles bound over a very broad area of the sidearm from the shoulder to the head (Figure 5A, lower) and an extraordinary form of the sidearm, probably not containing p150, was observed (Figure 5–S1A). Considering overexpression of p50 disrupts the sidearm from binding to the Arp1 rod (Echeverri et al, 1996), we presumed that modification of p50 C-terminus inhibited proper incorporation of p150 into the complex and disrupted sidearm formation. In contrast, p24-C-His exhibited typical morphology of sidearm, but showed no specific nanogold labeling (data not shown) probably due to steric hindrance because specific labeling was restored by exchanging sites of His-tag and SBP-tag (Figure 5–S1B). Collectively, the results suggest that p50 is situated in the shoulder domain with p24 and the C-terminal side of p50 might be associated with the C-region of p150, tethering it to the upper part of the shoulder.

**Figure 5.**
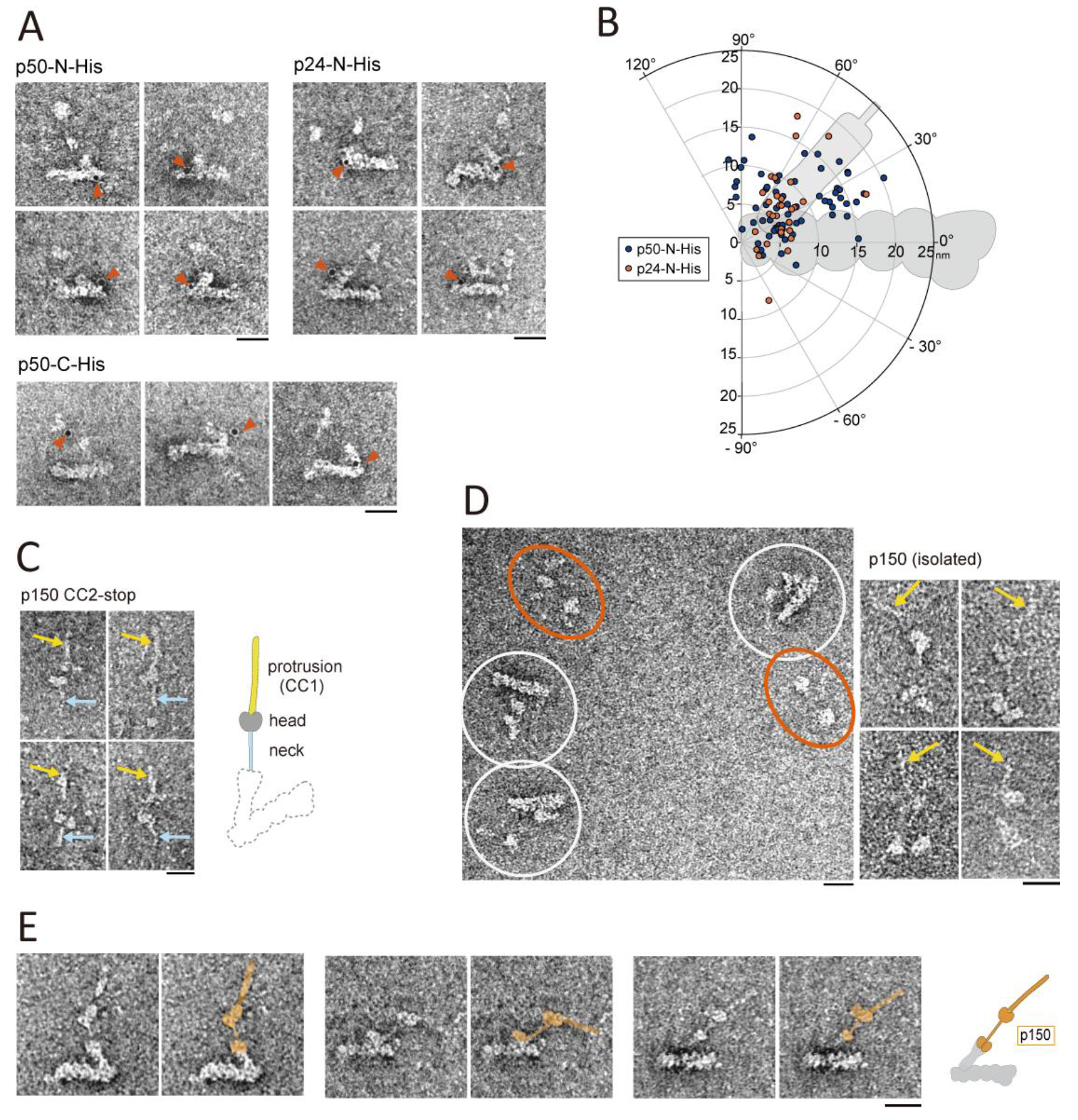
The shoulder domain is formed by p50, p24 and C-region of p150. (A) EM images of p50′N′His (upper left), p24-N-His (upper right) and p50-C-His (lower) labeled with the gold nanoparticles (red arrowheads). See also Figure 5—S1. (B) The distribution of the gold nanoparticles in p50-N-His (dark blue, *N*= 64) and in p24-N-His (orange, *N*= 32). (C) EM images of p150 CC2-stop, which lacks the C-region. The head, the protrusion formed by CC1 (yellow arrows) and the neck formed by CC2 (light blue arrows) are observed but the shoulder and the Arp1 rod are not, as illustrated in the cartoon on the right. (D) Left, a general view of the EM image of a mutant complex (p150-N-His). As well as the dynactin complex (white circles), isolated p150 particles (orange ellipses) are seen because the mutant p150 is more abundant than other subunits in cells exogenously expressing p150. Right, a gallery of EM images of isolated p150 dimers (p150-N-His) (E) Pairs of EM images of the dynactin complex (SBP-p62, containing a wild type sidearm) and identical ones with the supposed location of p150 pseudo-colored in orange. Bars represent 20 nm.

When the C-region of p150 was truncated (p150 CC2-stop in Figure 1–S1), neither Arp1 nor p50 was co-purified with this mutant (Figure 1–S2C) and EM observation revealed that no Arp1 rod was present and only a distal part of p150 (i.e., the sidearm without the shoulder) was found. These results demonstrated that the C-region of p150 constitutes a part of the shoulder domain and is indispensable for p150 incorporation into the dynactin complex via the remaining part of the shoulder composed of p50 and p24. This conclusion agrees with the recent cryo-EM study that revealed that several α-helices, assigned as the p150 C-terminus, p50 and p24, were bundled in the shoulder (Urnavicius et al., 2015). Disruption of structural integrity of the shoulder by p50-C-His (Figure 5–S1A) suggests that the C-terminus of p50 is a candidate for the dimerization domain of p50-p24 heterotrimers within the shoulder, which was visualized but not assigned in Urnavicius et al. (2015). Additionally, deletion analysis of p150 C-region (Figure 5C) may explain the rough eye phenotype of the *Glued^1^* mutation found in fruit flies (McGrail et al., 1995; Fan and Ready, 1997): transcription of p150 in *Glued^1^* is terminated in the middle of CC2 (Swaroop et al., 1985), which means C-region is absent, and consequently complex integrity of dynactin might be impaired in the mutant.

We found an additional evidence for further studying the incorporation of p150 into the dynactin complex. In the preparation of p150 mutants, isolated forms of p150 dimers were frequently observed with the whole dynactin complex (Figure 5D, left). Because p150 is more abundant than other components in cells exogenously expressing recombinant p150 (Figure 1–S2C), some p150 molecules should exist as isolated p150 dimers. Compared to the EM images of p150 CC2-stop (Figure 5C), the isolated p150 dimers had additional two oval structures (Figure 5D, right). This domain is likely to be composed of the C-region of p150 or perhaps might also include p50-p24 heterotrimers. Together with the deletion analysis of the C-region of p150, these two oval structures were considered to be essential for forming the shoulder domain and responsible for the complex formation. We also observed that gold nanoparticles bound to the His-tagged site of the isolated p150 dimer of each mutant (Figure 5–S2, red arrow heads), which agreed well with the results of the complex (Figure 3C, 4C). These isolated p150 dimers were similar in appearance to the shoulder/sidearm structures obtained by KI treatment and viewed by rotary shadowing (Eckley et al., 1999).

In the EM images of the whole dynactin complex, the most distal part of the shoulder was slightly swollen and divided into two smaller buds, which now we interpret to include the C-regions of p150 dimer. Considering these observations and the nanogold labeling of p150-C-His (Figure 2C,D), we presume that a dimer of p150 is situated in the dynactin complex as indicated in Figure 5E.

### Detailed structure of the head domain

The observation that CC2-start was localized at the head domain raised the possibility that the region between CC1 and CC2 constituted a part of the head domain. This region is rich in helix (Figure 6–S1A) and the cryo-EM study assigned this inter coiled-coil domain (ICD) as the head domain by its mass (Urnavicius et al., 2015). To confirm the possibility, we made deletion mutants lacking the entire or a part of ICD (Figure 1–S1, Figure 6–S1A). In the mutant lacking the entire ICD (p150 ΔICD), the head domain was almost never observed and the outline of the complex distal to the shoulder domain was quite obscure (Figure 6A, left; Figure 6–S1B). For mutants lacking the former or the latter half of the region (p150 ΔICD-a and p150 ΔICD-b, respectively; Figure 6A, middle and right), the head domains were more distinct than those of p150 ΔICD, but their head sizes appeared to be slightly smaller than that of the wild-type p150 (Figure 1A). In all of three ICD deletion mutants, the sidearm structure below the head, namely the neck and shoulder, remained unchanged compared with the wild-type sidearm. In contrast, CC1 was not distinctly observed except for p150 ΔICD-b (Figure 6A, right, yellow arrows). These results indicated that ICD is a part of the head domain and the former half (ICD-a) is important for the structural integrity of CC1.

**Figure 6.**
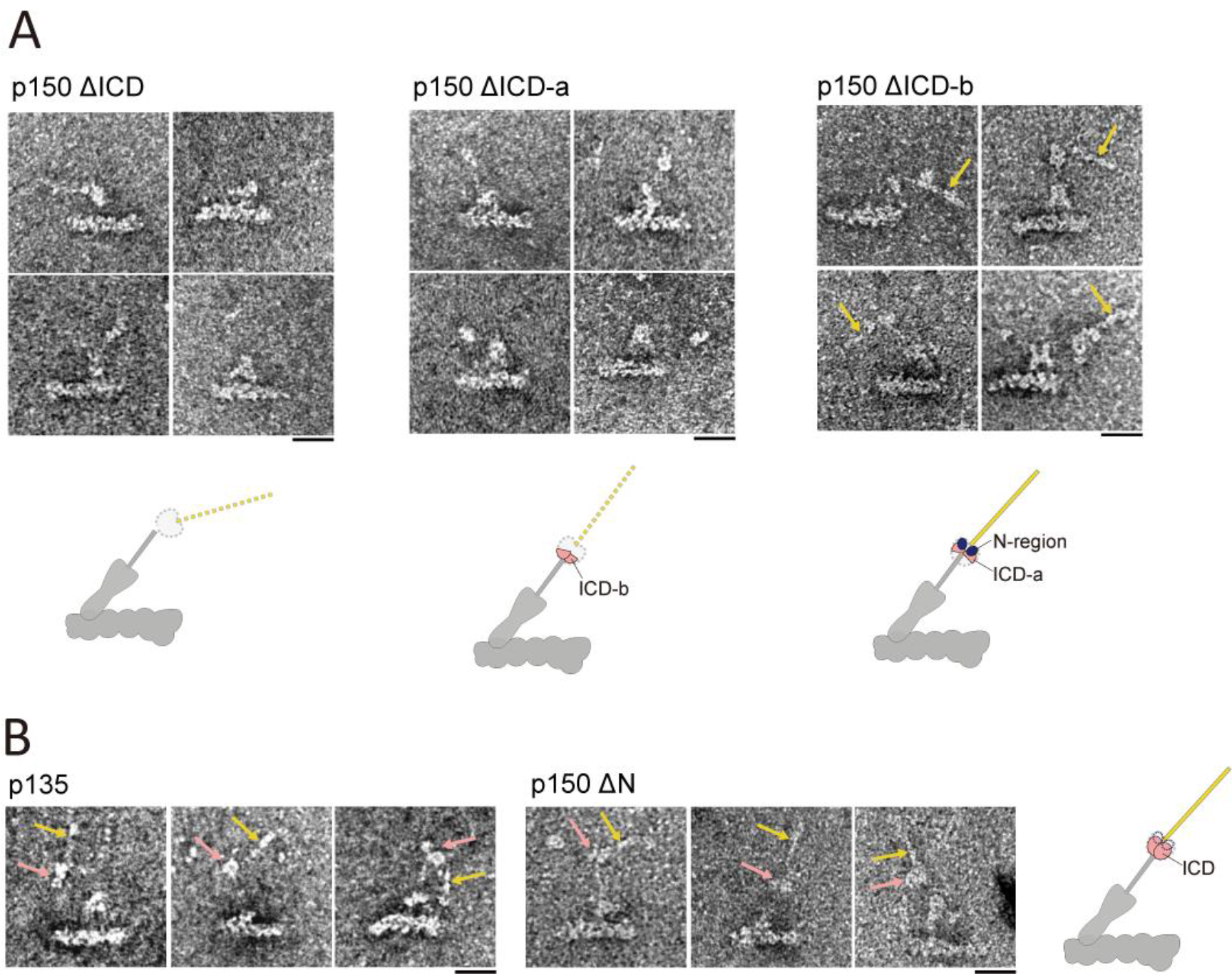
Identification of the p150 ICD. (A) EM images of p150 ΔICD (left), p150 ΔICD-a (middle) and p150 ΔICD-b (right). Yellow arrows indicate CC1. (B) EM images of the p150 mutants lacking the N-region: p135 (left) and p150 AN (right). Yellow arrows indicate CC1. Pink arrows indicate the head domain formed by ICD without the N-region.

Next, we investigated into the contribution of the N-region to the head domain. p135 is a splicing isoform of p150 expressed in mammalian neurons that lacks most of the N-region (Dixit et al., 2008; Tokito et al., 1996). Thus, we made two N-region deletion mutants: one mimicking p135 (p135) and the other lacking the entire N-region and starting at CC1 (p150 ΔN) (Figure 1–S1, S2C). Intriguingly, the morphology of these two mutants did not differ from that of the full length p150 and the sizes of the heads were indistinguishable from that of p150 (Figure 6B, pink arrows). Furthermore, CC1 of both p135 and p150 ΔN were observed to be similar to that of wild-type p150 (Figure 6B, yellow arrows). These results suggest that the contribution of the N-region to the size of the head domain may be considerably small when compared with that of ICD.

### CC1 adopts folded and extended forms

We have already demonstrated that ICD is an essential constituent of the head domain (Figure 6) and the N-region is located around ICD (Figure 2A,B). We next focused on how these two domains are structurally linked. CC1 lies between the N-region and ICD and it was occasionally observed to be unfolded (e.g., Figure 1A, panel g; Figure 4E, right panel) or more extended (Figure 7A) when compared with the ~ 30 nm protrusion (Figure 1–S3C,D). Here, CC1 was measured as being more than 50 nm in some molecules; however, the precise measurement was difficult because the size of the N-region (~ 190 aa) in wild-type p150 was too small for EM observation. Thus, we fused GFP to the N-terminus of p150 (p150-N-GFP in Figure 1–S1, S2C) expecting that the mass of the GFP + N-region (GFP-N) was sufficiently large and detectable in EM. We indeed observed that GFP-N (~ 490 aa) was as large as ICD (~ 470 aa) and both domains exhibited similar globular structures. We noticed that the distance between GFP-N and ICD was varied considerably (Figure 7B,C). When GFP and ICD were in close proximity, both structures situated at the base of CC1 (Figure 7B).

**Figure 7.**
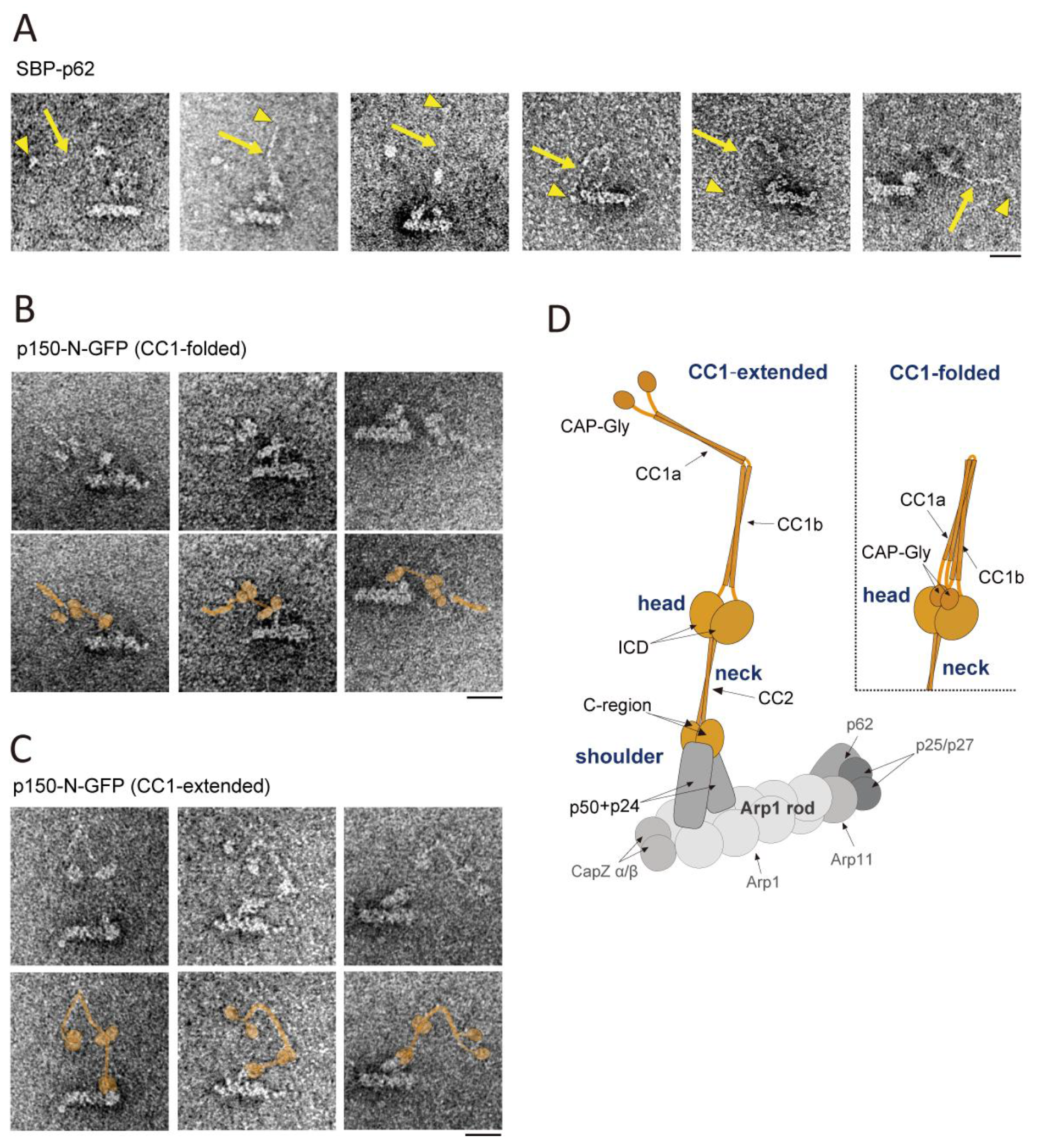
Conformational change of CC1 and new model of the dynactin sidearm. (A) EM images of a subset of SBP-p62 with longer CC1 (yellow arrows). Yellow arrowheads indicate N-region at the tip of CC1. (B), (C) Pairs of EM images of p150-N-GFP and identical ones with the supposed location of p150 pseudo-colored in orange. (B) ICD and the GFP + N-region are in proximity at the bottom of CC1 (CC1-folded). (C) ICD and the GFP + N-region are observed as separate and distinguishable structures, with these two domains bridged by a hooked CC1 (CC1-extended). (D) New model of the dynactin sidearm. The sidearm is composed of four domains: a protrusion, two heads, a neck and a shoulder. p150 is colored orange and each domain in p150 is indicated. CC1 takes a folded form (upper) and an extended form (lower). The model of the Arp1 rod and shoulder domains are based on the previous reports (Chowdhury et al., 2015; Eckley et al., 1999; Imai et al., 2006; Maier et al., 2008; Schroer, 2004; Urnavicius et al., 2015). Bars represent 20 nm.

In other cases, however, GFP-N was observed as a separate and distinct domain from ICD, with these two domains bridged by a hooked CC1 (Figure 7C). This finding indicates that CC1 adopts not only a folded form, which was reported in this and previous studies, but also an extended form, which was observed here for the first time. The co-existence of both forms of CC1 in a single sample suggests that CC1 undergoes a conformational change between the two forms. CC1 contains two adjacent parallel coiled-coils, CC1a and CC1b, joined by the hinge and it is likely that they contact each other and form a supercoil in the folded form, whereas they exist as two separate coiled-coils in the extended form. In line with this, CC1 was thinner in the extended form (Figure 7C) than in the folded form (Figure 7B). Varying visibility of CC1 described above (Figure 1A, panel a-h) may have been derived from the existence of this thinner form of CC1, i.e., CC1 may be more difficult to detect by EM in the extended form than in the folded form. When it was visible, CC1 in the extended form exhibited more various morphologies and was probably more flexible than in the folded form (Figure 7C, Figure 7–S1). Furthermore, owing to the GFP fusion, we also found that the two N-regions within the p150 dimer in the extended form were able to separate from each other (Figure 7C, middle and right; Figure 7–S1, type 4).

On the basis of our GFP fusion, nanogold labeling and deletion analyses of the p150 mutants, we present a new model for the domain organization of p150 and how it is situated in the dynactin complex via p50-p24 (Figure 7D). Since CC1 adopted either the folded or extended forms, we propose that CC1 acts like an “arm”, which undergoes a large conformational change (CC1-folded and CC1-extended in Figure 7D).

## Discussion

### Comparison of the sidearm structure by negative stain EM and cryo-EM

In this study, we investigated into the molecular architecture of the dynactin complex by negative stain EM. Whereas the length and thickness of the Arp1 rod is fairly constant, the orientation and configuration of the sidearm is remarkably diverse (Figure 1). Contrary to the “averaging” approach now popular in EM studies, we explore the structure of the sidearm exclusively by observation of individual molecules, to visualize even the most flexible part of the sidearm. Utilizing nanogold labeling and truncated mutants of the human dynactin complex, we revealed how p150 was folded and located within the distal side of the sidearm. Domain organization of p150 determined by our approach (summarized in Figure 7–S2) is generally consistent with the proposed interpretation of the cryo-EM image of pig brain dynactin (Urnavicius et al., 2015). Both models markedly differ from the previous model (Schroer, et al., 2004) especially in assignment of CC1 as the protrusion from the head and ICD as the head.

Despite their general agreement in the sidearm organization, our results exhibit some important differences from the cryo-EM results (Urnavicius et al., 2015). The first is visibility of the N-terminus of p150. We found that the N-region located either in close proximity to or far from the head (Figure 2, Figure 7); whereas, in their cryo-EM, the N-region disappeared and the N-terminus of CC1a was unclear probably because the p150 N-terminus was extremely flexible and their sample included the isoform lacking the N-region (p135) as well as the full length p150. The second is orientation and conformation of CC1. In the class average of their cryo-EM, CC1 was visualized only in the folded form and was always docked to the Arp1 rod. In our observations, however, CC1 adopted a wide range of orientations with respect to the neck direction (Figure 3C,D) and exhibited the folded and extended forms (Figure 7, Figure 7–S1). The conformational change in CC1 adds further morphological variations to the sidearm, which might facilitate its interaction with dynein and MTs. As to the discrepancy with the cryo-EM results, we found that the majority of the sidearm population was in the undocked form under our condition (Figure 1) and the docked particles might be preferred in the process of single particle analysis, like shown in fig. S7 in Urnavicius et al. (2015).

The third is flexibility of the shoulder. In our results, the distal end of the shoulder (the distal shoulder) also moved flexibly (Figure 1B, green plots; Figure 4C,D) and it clearly detached from the Arp1 rod in some molecules (e.g. Figure 5E, left and right panels); the distal shoulder was more rigid and closer to the Arp1 rod in their cryo-EM or class averages from other groups (Imai et al., 2006; Imai et al., 2014, Chowdhury et al., 2015). This apparent contradiction might also come from the difference between non-averaged or averaged dynactin images rather than difference among the samples because the distal shoulder was also observed to be flexible and detached from the Arp1 rod in non-averaged images of chick embryo dynactin (Schafer et al., 1994; Imai et al., 2014). Our mutant analysis and the cryo-EM results on the shoulder domain (Figure 5; Urnavicius et al., 2015) indicate that p150 is incorporated into the complex by interaction of the C-region of p150 and the two p50-p24 arms. Thus, the flexibility of the shoulder observed in our non-averaged dynactin suggests that although the p50-p24 arms looked firmly attached to the Arp1 rod in averaged images, they have the potential to move dynamically and regulate the orientation of the sidearm including p150 (Figure 1–S4).

Summing up, the differences between our and previous cryo-EM results mostly come from the fact that we focused on the undocked and flexible sidearm with diverse structures whereas previous cryo-EM studies preferred the docked and rigid sidearm with a uniform structure. We suppose that structural analysis based on averaging technique alone is not sufficient to fully comprehend the nature of flexible proteins like dynactin sidearm and that non-averaging approach is also needed to decipher its functional structure. Whether dynactin in the cell adopts both docked and undocked forms to play distinct physiological roles is a next intriguing question.

### Labeling of flexible objects by gold nanoparticles

Chemically modified gold nanoparticles visualize the tagged sites within protein complexes (Ichikawa et al., 2015; Guesdon et al., 2016; Song et al., 2015). Using gold nanoparticles with diameter less than 5 nm (Kitai et al. 2011), the specific sites within ~30 nm CC1-protrusion (Figure 3, Figure 3–S1) and ~20 nm CC2-neck (Figure 4) were clearly discernible. This provided sufficient information to determine the folding pattern and the domain organization of p150 in the sidearm through the observation of individual dynactin molecules.

Application of nanogold labeling is not confined to rigid or static structures. For example, it was used to label microtubules exhibiting dynamic instability (Guesdon et al., 2016). In the present study, we labeled numerous sites along the sidearm (Figure 7–S3, Table S2). The distributions of gold nanoparticles seen in Figure 7–S3A probably reflected the flexibility of the sidearm, and were consistent with the measurement of the position of the head and base of the neck without labeling (Figure 1B, right). Thus, we consider the technique as advantageous in deciphering the structure and conformation of flexible objects like the dynactin sidearm without compromising its flexibility and morphological heterogeneity.

### CC1 as a possible regulator for interaction with dynein and MT

There are some descriptions of the dynein-binding region within the CC1 region (Siglin et al., 2013; Tripathy et al., 2014). Moreover, binding of CC1 with dynein was shown to have positive and negative effects on dynein motility, depending on the length of the fragment or the existence of adjacent domains (Kobayashi et al., 2017; Tripathy et al., 2014). Contrary to the well-known inhibitory effect of the CC1 fragment on dynein function in cells (Quintyne et al., 1999) or in cell extract systems (Ishihara et al., 2014; Suzuki et al., 2017), its mechanism of action and *in vitro* effect on dynein motility remains controversial as described in Introduction. In this study, we found that CC1 folded back at the CC1-hinge (Figure 3), and that CC1a and CC1b had contact with each other (folded form) or separated (extended form) (Figure 7). We speculate that the affinity of CC1 toward dynein IC and its apparently ambiguous effect on dynein motility are regulated by the conformational change of CC1. For example, an intramolecular interaction between CC1a and CC1b may regulate its effect on dynein, as proposed in the previous study (Tripathy et al., 2014). Moreover, it was recently discovered that dynein tail interacts with the Arp1 rod with the aid of cofactors (Chowdhury et al., 2015; Urnavicius et al., 2015) and a functional relationship between this mode of interaction and the IC-CC1 based interaction is an open question.

In mammalian cells, the MT binding property of the dynactin is controlled by expression of several isoforms in which alternative splicing affects the composition of the N-region (Dixit et al., 2008; Zhapparova et al., 2009). The p150 constructs used in this study (p150-B or DCTN1B) lacked exon 5-7 (basic-rich domain). We examined previously MT-binding ability of this isoform by TIRF microscopy and found that isolated p150-B rarely bound to MTs *in vitro* but deletion of CC1 domain remarkably restored the intrinsic MT-binding ability of CAP-Gly (Kobayashi et al., 2006; Kobayashi et al., 2017). The conformational change of CC1 revealed by the present study offers a simple explanation for the difference in MT-binding ability. In this model, N-region including CAP-Gly is usually masked by the folded form of CC1 and this state inhibits the binding of the N-region to MTs; however, if the whole CC1 domain is deleted or CC1 is turned into the extended form, the N-region is un-masked and binds to MTs.

Previous studies showed that the interaction between CAP-Gly and MTs was not always required for processive movement of dynein (Kardon et al., 2009) but was necessary for the initiation of dynein-driven retrograde transport of vesicles from the neurite tip (Lloyd et al., 2012; Moughamian et al., 2012) and for the initiation of ultra-processive movement of DDB (dynein, dynactin and BICD2) complex at tyrosinated MTs (McKenney et al., 2016). Regulation of CAP-Gly by CC1 may be important for understanding these differential effects of CAP-Gly on MT-based transport. In another cellular context, the regulation of dynactin MT-binding ability may play a key role in a passive transportation of dynein to the MT plus ends by +TIPs (Akhmanova and Steinmetz, 2008), i.e., how dynein molecules are allotted to two pools of active motility and passive transportation examined in reconstitution assays (Baumbach et al., 2017; Jha et al., 2017) is possibly linked with the conformation of CC1.

This study reveals the flexibility of the sidearm, especially that of CC1, which is likely to have functional meaning. However, all regulatory roles of CC1 proposed here are based on the premise that the conformational change of CC1 actually happens and is controlled in the cell. Thus, determination of the conformation of CC1 *in vivo* and validation of conformational transition *in vitro* are the next important challenges.

## Materials and methods

### Construction of expression vectors

cDNAs encoding the dynactin subunits were amplified from HEK-293 cells by RT-PCR. The PCR primers used for cloning of dynactin subunits are as follows: 5′-ATGGCACAGAGCAAGAGGCAC-3′ and 5′-TTAGGAGAT GAGGCGACTGTG-3′ for p150 (DCTN1, NM_004082), 5′-ATGGCGTCCTTGCTGCAGTCG-3′ and 5′-TTAAGGAAGAAGTGGGCCCAA-3′ for p62 (DYNC4, NM_001135643), 5′-ATGGCGGACCCTAAATACGCC-3′ and 5′-TCACTTTCCCAGCTTCTTCAT-3′ for p50 (DCTN2, NM_006400), and 5′-ATGGCGGGT CT GACT GACTTG-3′ and 5′-TCACTCCTCTGCTGGCTTCAC-3′ for p24 (DCTN3, NM_007234). Note that our cloned dynactin sample of p150 from HEK-293 cells was DCTN1B isoform, which lacked some amino acids of the basic-rich domain (Δexon 5-7) as reported (Dixit et al, 2008 and Zhapparova et al, 2009). We found a missense mutation (c.2566G>C) which produces p.A856T in the ICD region; however, we confirmed that the mutation does not affect the dynactin structure. The primers used for construction of p150 mutants are summarized in Table S3. For p135 construction, N-terminal 131 amino acids were deleted and 17 amino acids were added to its N-terminus to mimic p135 isoform found in neuron (Tokito et al, 1996).

The PCR products were cloned into pcDNA5/FRT/TO (LifeTechnologies, Carlsbad, California), a tetracycline-inducible mammalian expression vector. For protein purification, the streptavidin-binding peptide (SBP) tag was fused at either the N- or C-terminus of the dynactin subunits. For Ni-NTA Au nanoparticle labeling, an octa-histidine tag (His-tag) was inserted in either the N- or C-terminus or internally through a short linker (Gly-Gly-Gly-Ser) (see Figure 1–S1). The mutant p150-N-GFP has GFP fused at N-terminus of p150 as previously described (Kobayashi et al., 2017). A His-tag and SBP tag were included in the GFP and used for purification (Kobayashi et al., 2008). The insertion of these tags or deletion of specific domains was achieved by inverse PCR.

### Generation of the stable inducible HEK-293 cell lines

Flp-In T-REx HEK-293 cells (LifeTechnologies) were maintained in Dulbecco’s modified Eagle’s media (DMEM, LifeTechnologies) supplemented with 10% fetal calf serum and 2 mM L-glutamine. The stable inducible cell lines were generated by co-transfection of the pcDNA5/FRT/TO vector containing recombinant dynactin subunits with the pOG44 vector encoding Flp recombinase according to the manufacturer’s instructions. The transfectants were screened by including 100 μg/ml hygromycin and the hygromycin-resistant colonies were harvested.

### Purification of the dynactin complex

Purification of the dynactin complex was carried out using a SBP-tag (Ichikawa et al, 2011; Kobayashi and Murayama 2009). Stable HEK-293 cells were cultured in five 150-mm tissue culture dishes and protein expression was induced by doxycycline (2 μg/ml) for 48 h. Cells were harvested, washed twice with phosphate buffered saline, and homogenized in buffer A (50 mM Tris-HCl, pH 7.5, 0.15 M NaCl, 10% sucrose, 5 mM MgSO4, and 1 mM dithiothreitol) containing 0.5 mM ATP, 0.05% Triton X-100, and complete mini protease inhibitor cocktail (Roche, Basel, Switzerland). The lysate was centrifuged and the resultant supernatant was applied onto a StrepTrap HP column (1 ml) (GE Healthcare, Chicago, Illinois) pre-equilibrated with buffer A. After extensive washing of the column with buffer A, the bound proteins were eluted in buffer A containing 2.5 mM desthiobiotin. The fraction from the peak of interest was quickly frozen in liquid nitrogen and stored at −80 °C until use for EM observation.

### Mass spectrometric analysis to identify proteins

Peptide mapping was carried out using the Triple TOF 5600 mass spectrometer systems, which consisted of nano-ESI and TOF (AB SCIEX MA, USA). The Triple TOF 5600 mass spectrometer was combined with Eksigent NanoLC-Ultra system plus cHiPLC-nanoflex system (AB SCIEX MA, Framingham, Massachusetts) with attached 75 μm (id) × 15 cm Chrom XP C18-CL column. The solvent system consisted of (A) 0.1% formic acid/2% acetonitrile and (B) 0.1% formic acid/90% acetonitrile. The solvent was linearly changed from 2% B to 40% B for 40 min and subsequently fixed at 90% B for 5 min. The flow rate was 300 nl/min. Identification of proteins was performed using Protein Pilot 4.0 software (AB SCIEX MA) (Hayashi et al., 2013).

The proteins were subjected to digestion as described previously (Fujimura et al., 2008). Finally, the residue was dissolved in 30 μl of 0.1% formic acid. Aliquots were used for peptide identification by mass spectrometry.

### Negative stain electron microscopy and nanogold labeling

Samples were applied to pre-hydrolyzed carbon-coated EM grids and negatively stained with 1.4% (w/v) uranyl acetate solution, observed at 40,000× magnification in a transmission electron microscope, H7500 (Hitachi High-Techonologies, Tokyo, Japan) operating at 80 kV. Micrographs were taken using a 1024 × 1024 pixel CCD camera, Fast Scan-F114 (TVIPS, Gauting, Germany). The nominal magnification was 40,000×, giving a sampling of 2.6 Å/pixel. All experiments were performed independently at least two times. For each mutant, several EM grids were prepared and observed, and subsequently data were collected from the most adequately-stained grid. The images were not inversed for the figure presentation, except panels (a), (b), (e) and (f) in Figure 1.

Ni-NTA-gold nanoparticles were synthesized according to Kitai et al, 2011 using a modified method to reduce the diameter to 3 nm. The purified proteins were mixed with Ni-NTA-gold nanoparticles and incubated on ice for 30 min. The dynactin complex and Ni-NTA-gold nanoparticles was mixed to yield 10-20% labeled particles for all samples examined. These conditions were chosen to avoid non-specific binding of Ni-NTA Au nanoparticles. It is noted that the free Ni-NTA-gold nanoparticles rarely bound to the carbon surface of the grid, probably because of an electrostatic repulsion.

The length and angle in the EM images were measured using ImageJ software (NIH). Reference points of dynactin (Figure 1–S3A) and the center of the gold nanoparticles were determined by visual inspection. We used two different polar coordinates to measure and map these points (Figure 1–S3A, right). For the gold nanoparticles bound around the head of dynactin (Figures. 1D, 2D and 3B), the distance from the origin (X: the center of the heads) and the angle from the neck direction (XY) were measured and mapped as shown in Figure 1–S3A, right, red graph. For the gold nanoparticles bound along the shoulder of dynactin (Figures. 1F, 3D and 4B) and for the points within the sidearm (Figure 1B), the distance from the origin (P: the barbed-end of the Arp1 rod) and the angle from the Arp1 rod direction (PQ) were measured and mapped as shown in Figure 1–S3A, right, blue graph.

## Acknowledgments

This work was supported by JSPS KAKENHI Grant Numbers 25117503 and 25291031(to Y.Y.T.), 24590330 (to T.M.), 23300128 (to T.S.) and by MEXT-Supported Program for the Strategic Research Foundation at Private Universities, 2011-2015.

## Author Contributions

KSa, TM, TS and YYT designed the research. TM, TK and KSh made the expression vectors and purified the proteins, SK and TF carried out mass spectrometry analysis. KSa and TH performed EM, gold labeling experiments and the analysis. KSa, TM and YYT wrote the manuscript.

**Abbreviations List**
EM: electron microscopy
CC1: coiled coil 1
CC2: coiled coil 2
SBP: streptavidin-binding peptide
His-tag: octa-histidine tag

---

**Figure 1–S1.**
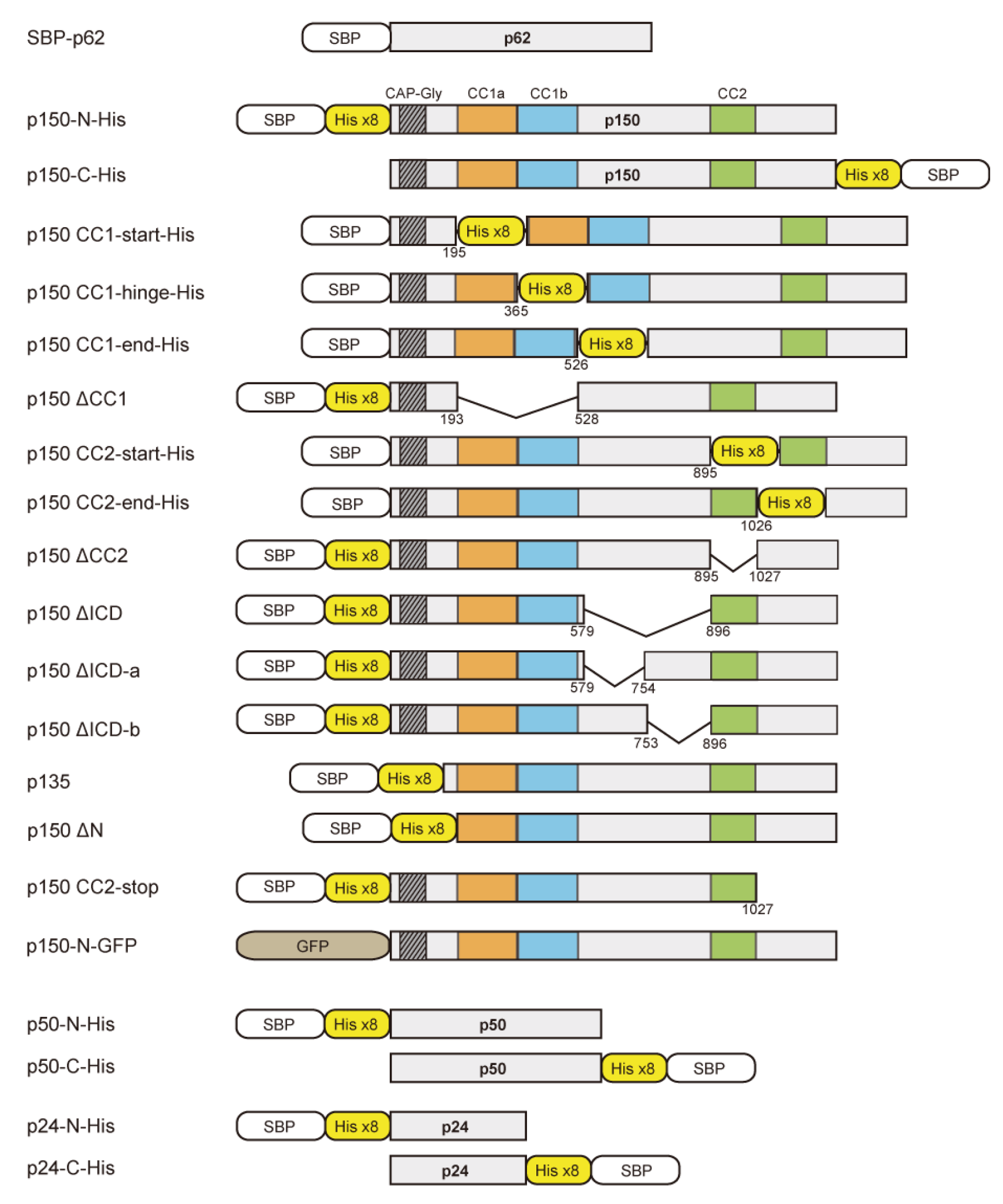
Constructs used in this study. The octa-histidine tag (His-tag) flanked by short linkers (GGGSHHHHHHHHGGGS) was inserted after K195, K365, Q526, E895, and E1026 for CC1-start-His, CC1-hinge-His, CC1-end-His, CC2-start-His, and CC2-end-His, respectively. For the other mutants, the His-tag was fused at either the N- or C-terminus. The residues S194-Q527, R896-E1026, G580-E895, G580-G753, and G754-E895 were deleted in ΔCC1, ΔCC2, ΔICD, ΔICD-a, and ΔICD-b, respectively. The C-terminal region encompassing L1028-S1253 was truncated in CC2-stop. Note that a His-tag and a SBP-tag were included in the GFP of the p150-N-GFP construct (see **Materials and methods**).

**Figure 1–S2.**
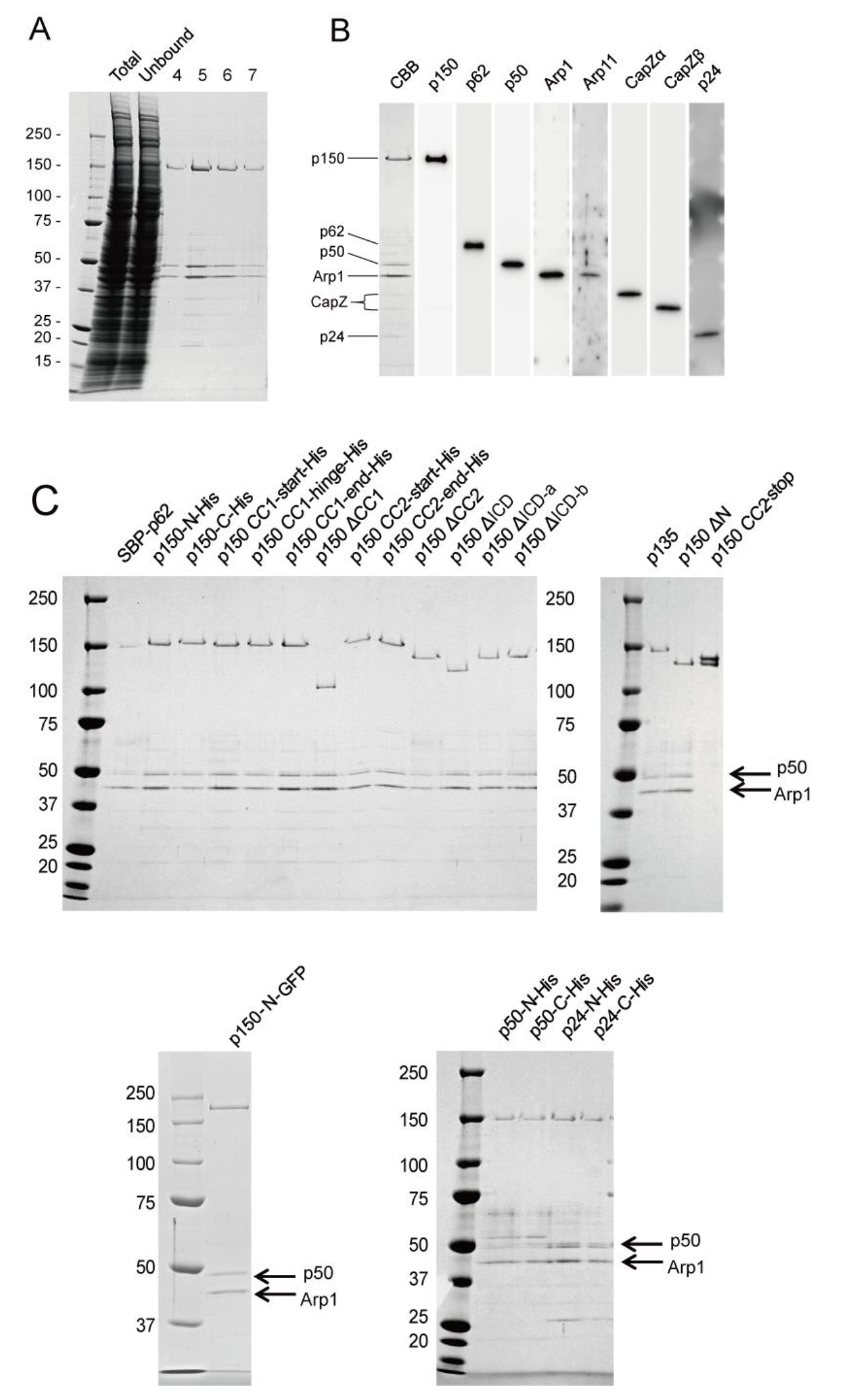
Sample preparation of recombinant human dynactin. (A) Purification of the dynactin complex by StrepTrap column chromatography. SDS-polyacrylamide gel (3%—15%) of the lysate from HEK-293 cells expressing p150-N-His (Total), the unbound fraction from the column (Unbound), and the eluted fractions No. 4—7. (B) Western blotting of the purified p150-N-His (No. 5 fraction in (A)). The purified fraction contained all the dynactin subunits examined (p150, p62, p50, Arp1, Arpl1, CapZα, CapZβ, and p24). (C) SDS-polyacrylamide gels of the purified dynactin mutants used in this study. p50 and Arp1 bands (arrows) were clearly detected in all the mutants except for p150 CC2-stop. Lower bands in p150 CC2-stop may be a degradation product.

**Figure 1–S3.**
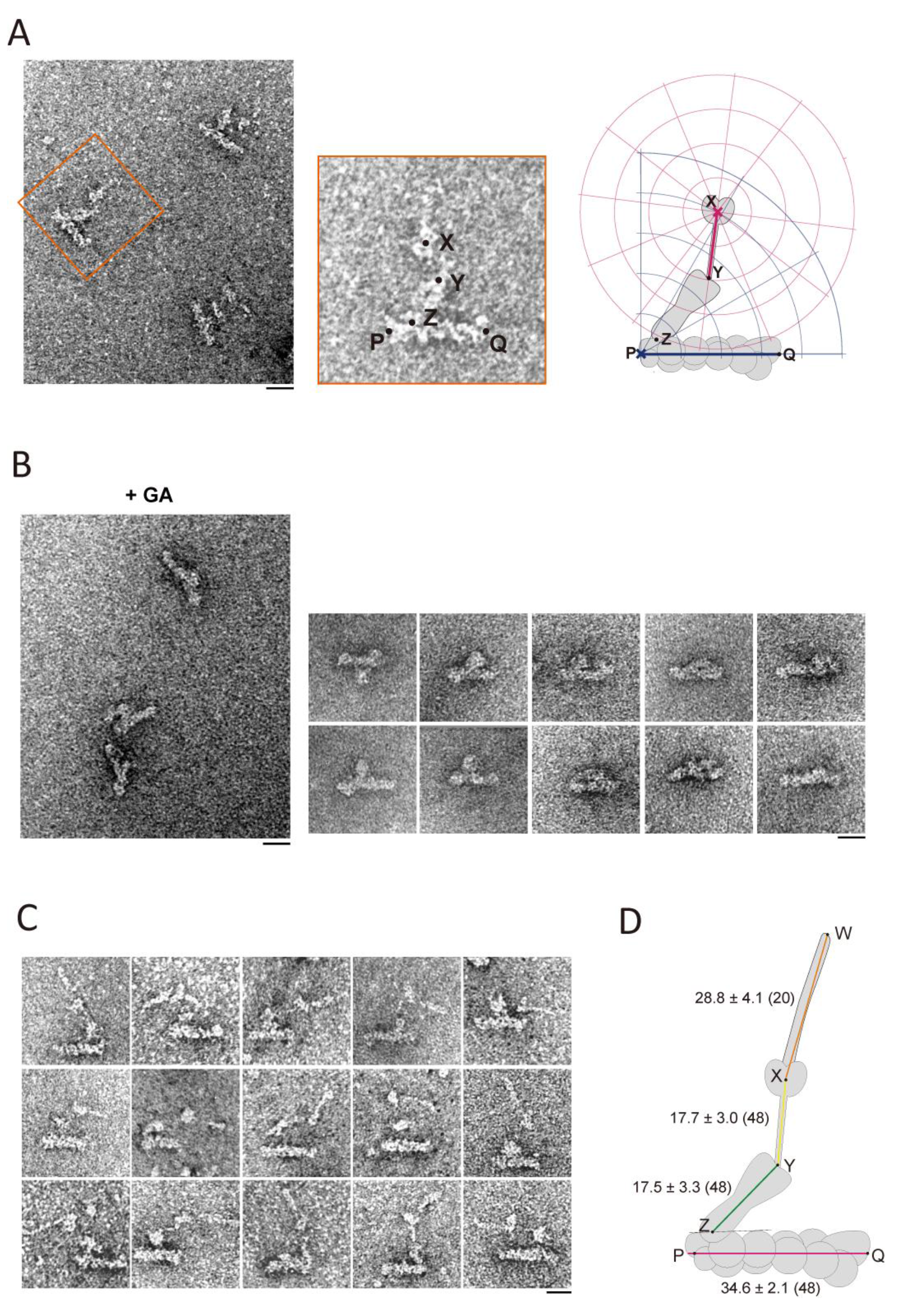
Morphological features of the dynactin complex and the effect of glutaraldehyde fixation. (A) sMorphological features of the dynactin complex. Left, a general view of the negative stain EM image of the dynactin complex (SBP-p62). Middle, a magnified image of the molecule designated by the orange box in the left general view. Points are assigned to show the morphological features of the dynactin complex and to define the reference points in it. P: barbed end of the Arp1 rod; Q: pointed end of the Arp1 rod; X: center of the two heads; Y: the neck-shoulder junction (base of neck); Z: proximal end of the shoulder. Right, illustration of how coordinate systems are set to measure the reference points in dynactin complex or positions of gold nanoparticles. See Materials and methods for details. (B) Glutaraldehyde fixation of the dynactin complex. Left, a general view; right, a gallery of molecules with the docked sidearm. The dynactin complex (SBP-p62), which includes wild type p150, was fixed with 2 % glutaraldehyde for 30 min on ice and then negatively stained. In most molecules, the shoulder appeared to closely contact with the Arp1 rod, and the neck, the head and the CC1-arm were not able to be recognized as distnguishable structures. Lower concentrations of glutaraldehyde(0.5 %, 1 %) and shorter periods of fixation time (5 min, 10 min) did not alter the result. Bars represent 20 nm. (C) A gallery of EM images of SBP-p62 containing a wild-type sidearm. Molecules with the filamentous structure were selected here (corresponding to Figure 1A, (e)—(h)). (D) Somatometry of the dynactin complex. The average lengths (mean ± SD (N) (nm) between the reference points are indicated. The definition of the reference points are the same as Figure 1–S3A, except for the tip of the filament (W). The data set is the same as the dynactin molecules with the undocked form in Figure 1B (*N*= 48). The data set includes a subset of molecules in which the tip of the filament was specified (*N*= 20).

**Figure 3–S1.**
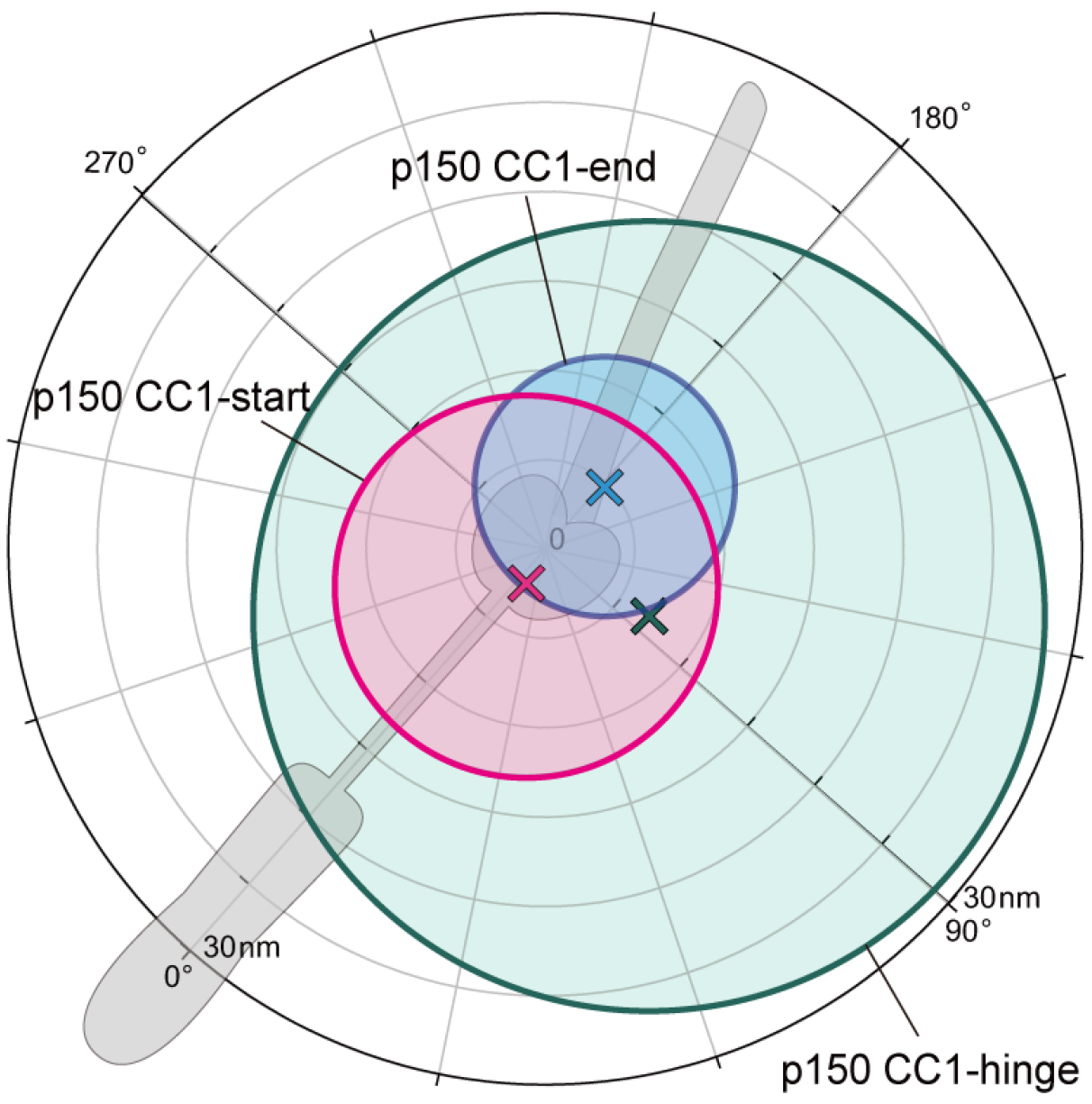
Distribution map of the nanogold-labeled sites in p150 CC1. The range of distributions of gold nanoparticles for p150 CC1-start-His (pink), p150 CC1-hinge-His (green) and p150 CC1-end-His (light blue). The crosses indicate the centroids. SDs of the nanogold distributions, calculated by root mean squared distance, are shown as radii of circles around the centroids (listed in Table S2). Original plot data were shown in Figure 3B.

**Figure 3–S2.**
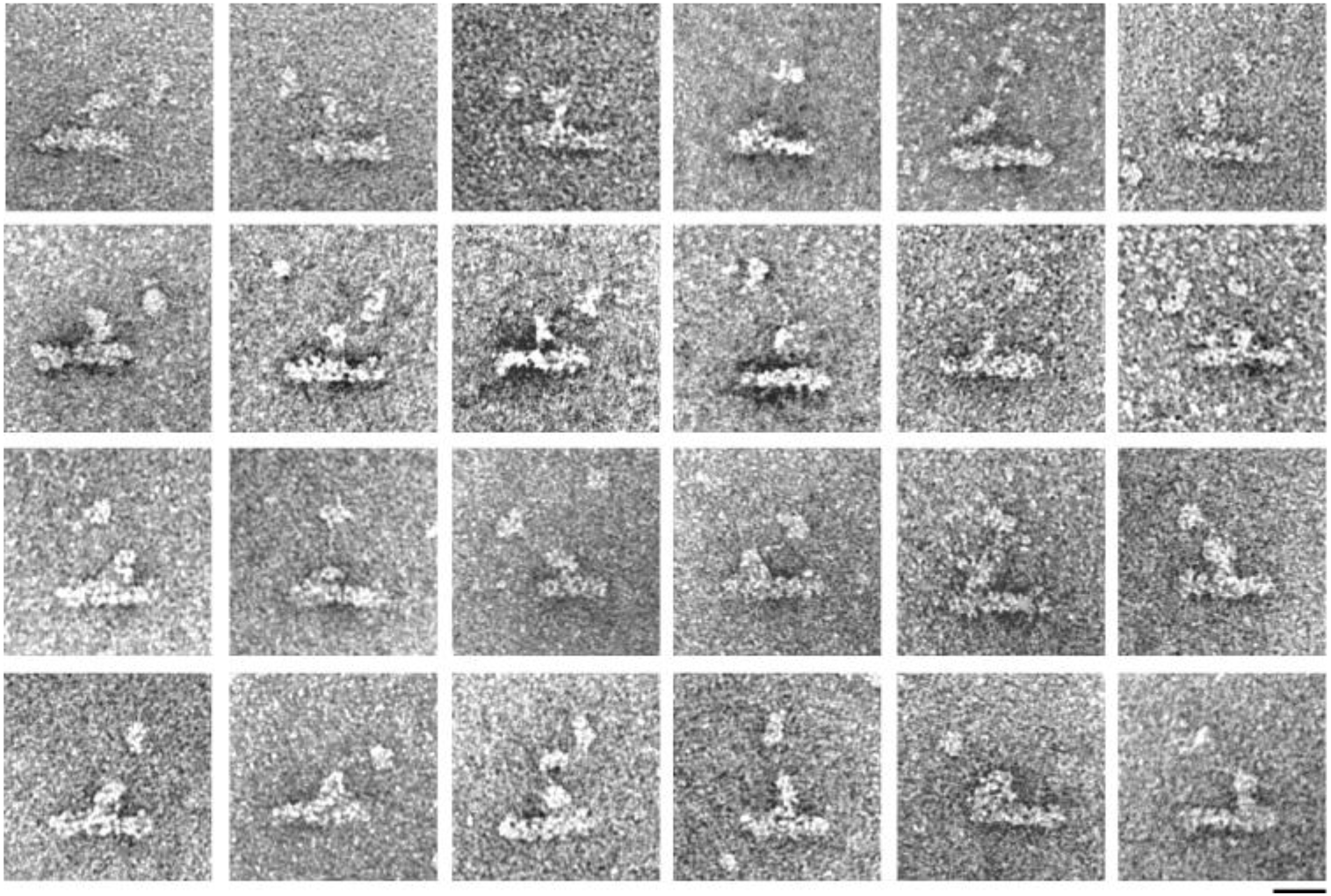
EM images of p150 ACC1. A gallery of EM images of p150 ACC1. The filament seen in Figure 1–S3C disappears in this mutant. Bar represents 20 nm.

**Figure 5–S1.**
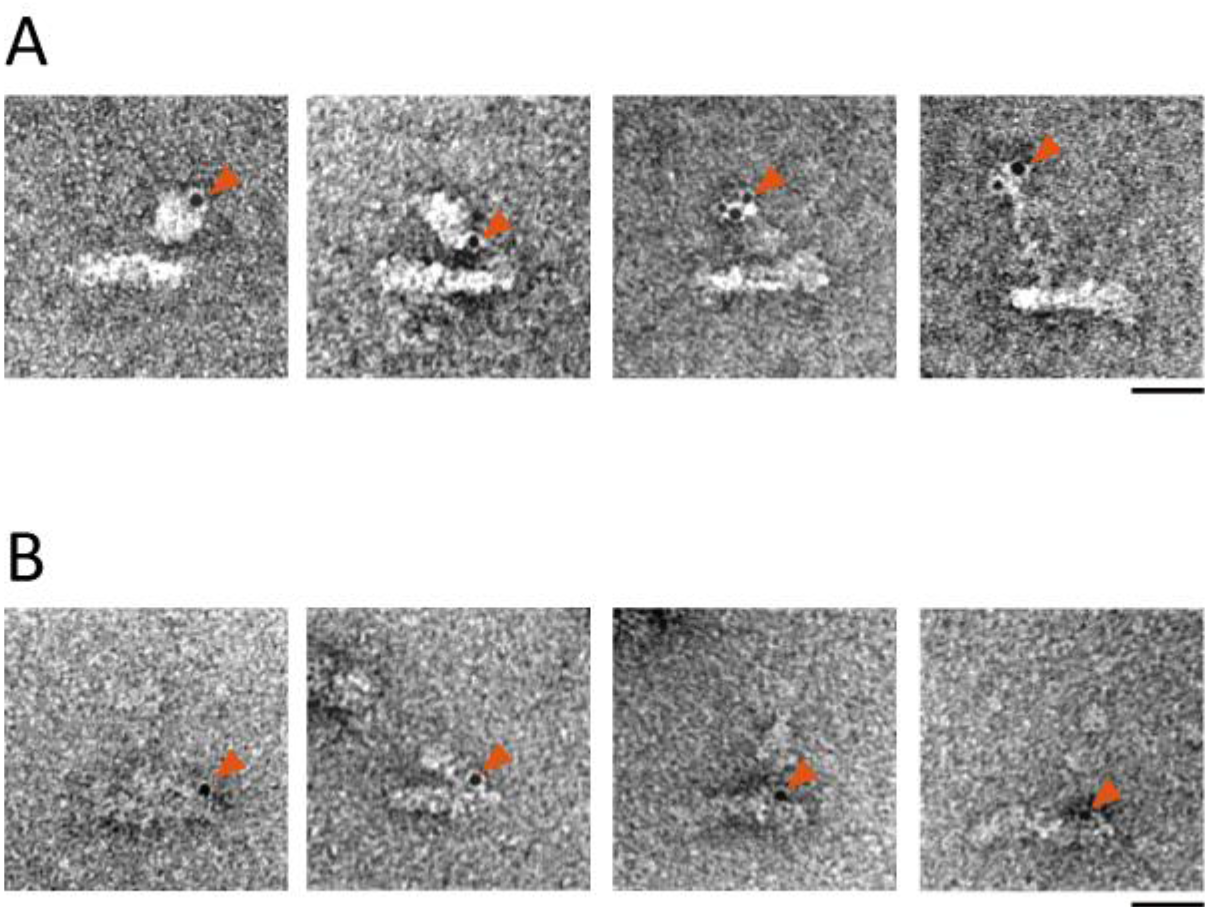
EM images and gold labeling of p50-C-His and p24-C-His mutants. (A) The dynactin complex of p50-C-His showing an irregular form of the sidearm. The sidearm in this mutant exhibited bigger and more expanded appearance than those of other mutants, in which the components of the sidearm might be misfolded (left two panels). In another case, a thin structure, which might be an unfolded form of p50, was observed to extend from the Arp1 rod (rightmost panel). Bound gold particles (red arrowheads) show the site of the C-terminus of p50 in each molecule. (B) The dynactin complex of the p24 mutant to which SBP and His tags are added to the C terminus of p24 in this order. Thus, the gold particles indicate the position of the SBP C-terminus. Bars represent 20 nm.

**Figure 5–S2.**
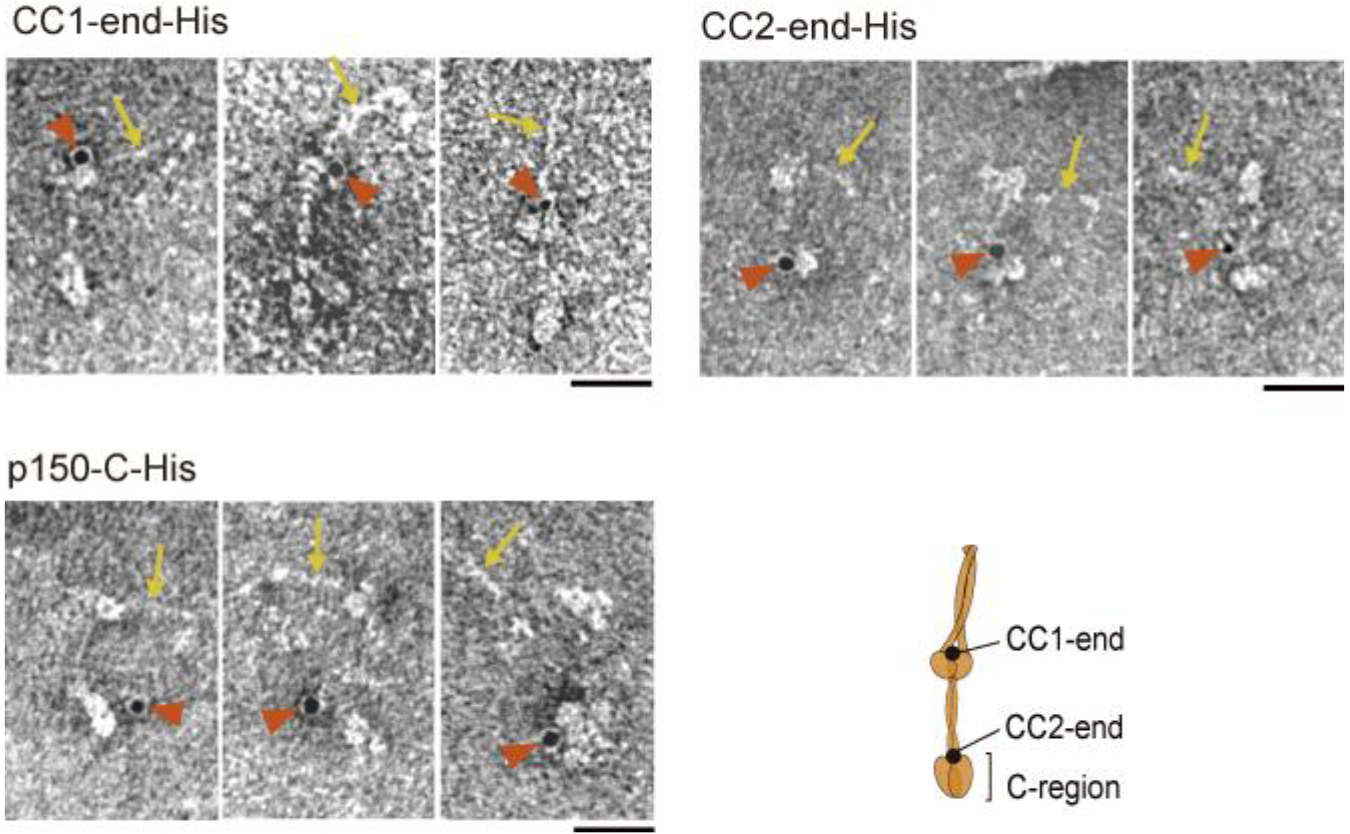
EM images and gold labeling of p50-C-His and p24-C-His mutants. EM images of isolated p150 dimers that are not incorporated into the dynactin complex in the mutants of p150: p150 CC1-end-His (upper left), p150 CC2-end-His (upper right), p150-C-His (lower left). Gold nanoparticles (red arrowheads) are bound to their His-tag sites and CC1 (yellow arrows) are observed.

**Figure 6–S1.**
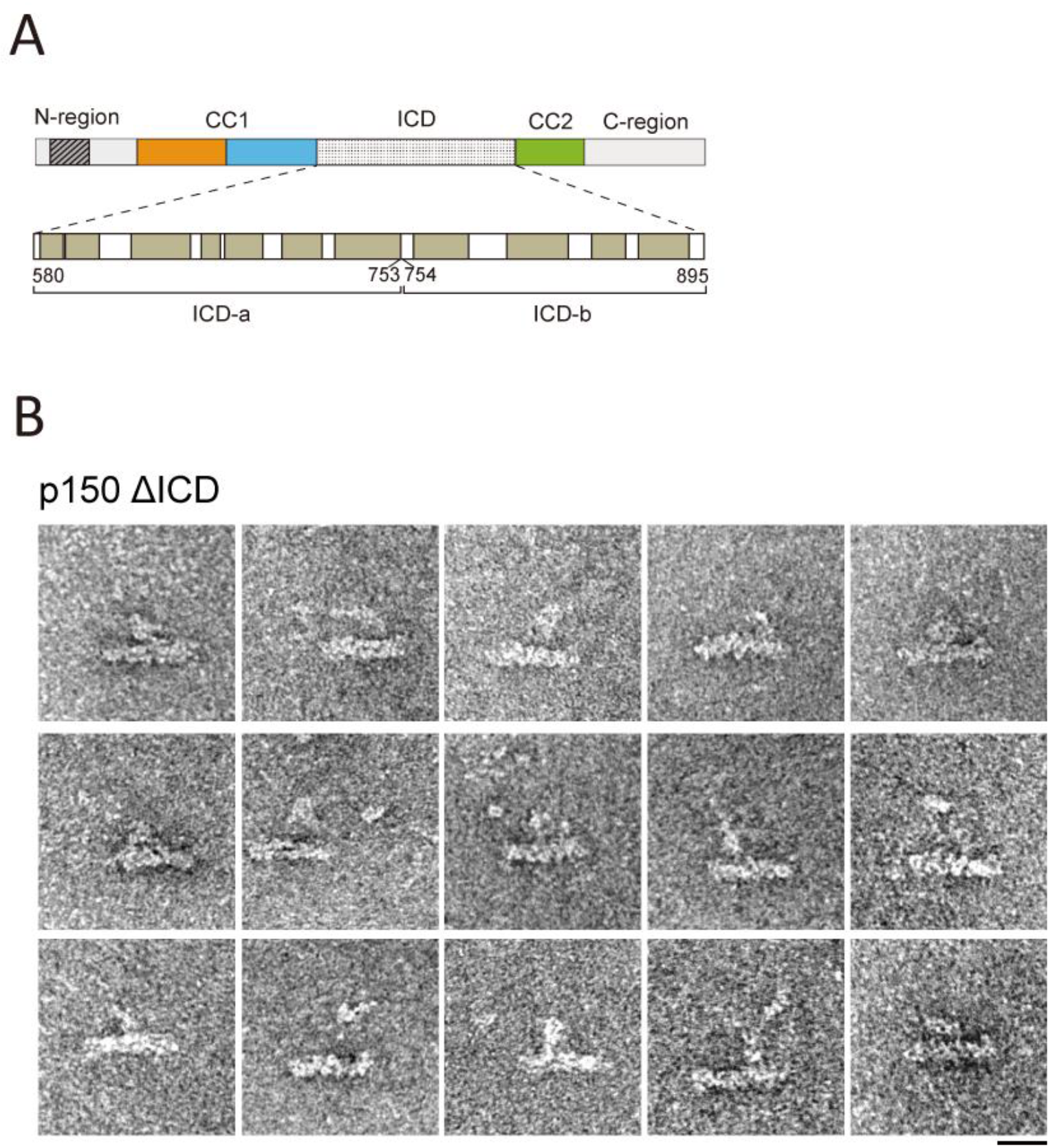
Structural contribution of ICD to the head. (A) Secondary structure prediction of ICD. The ochre gray colored regions of the sequence are predicted to form helices by PSIPRED (McGuffin et al., 2000). (B) A gallery of EM images of p150 ΔICD. Bar represents 20 nm.

**Figure 7–S1.**
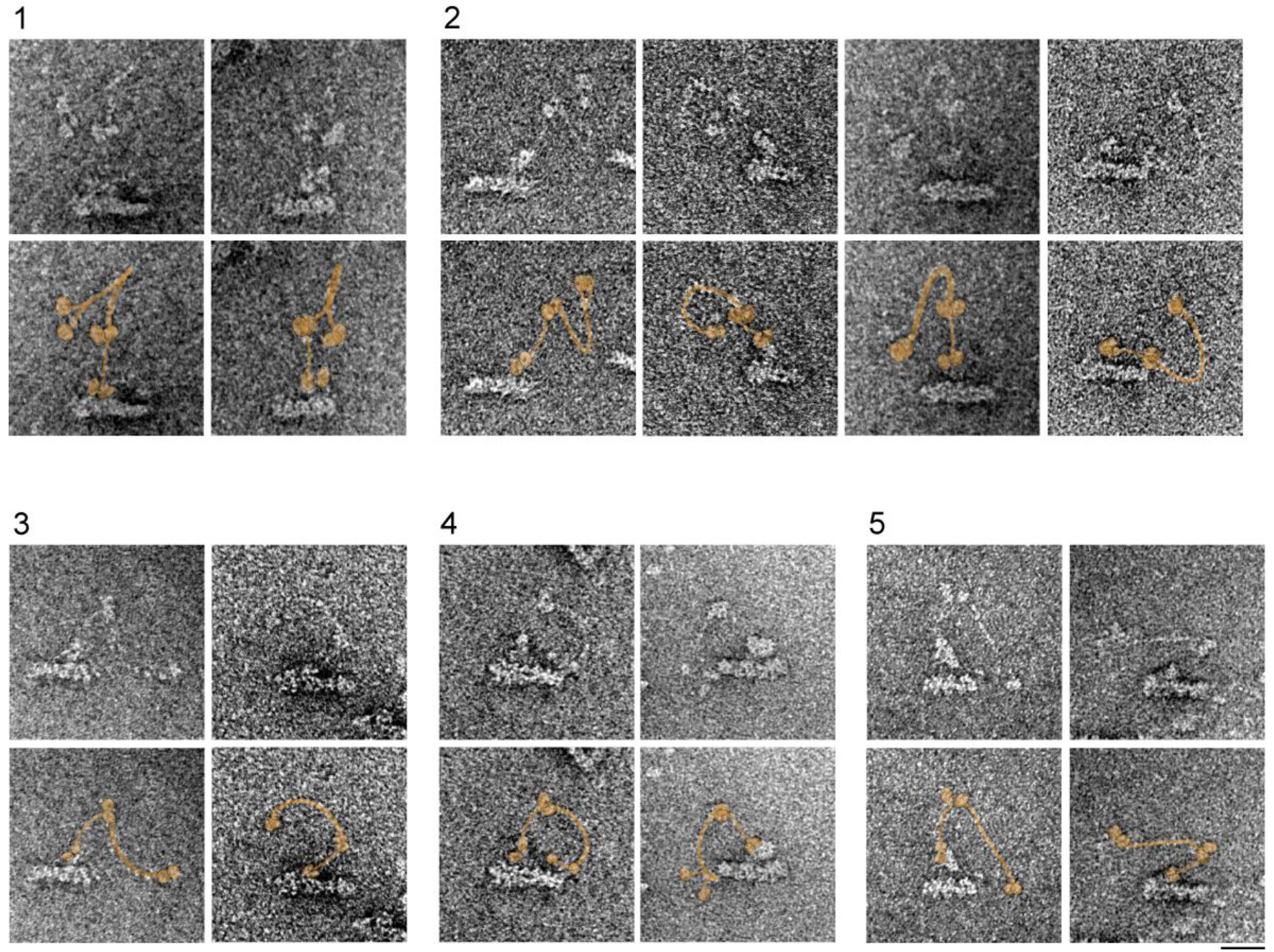
A gallery of the extended form of CC1. Pairs of EM images of p150-N-GFP and identical ones with the supposed location of p150 pseudo-colored in orange. Molecules with the extended (or not folded) form of CC1 are selected here and various morphologies are seen. We classified them as several morphological types. Type 1: CC1a and CC1b partially contact each other in the region near the hinge and ICD and N-GFP are separated; Type 2: CC1a and CC1b are separated and the angle made by ICD, the hinge (the middle of CC1) and N-GFP is acute; Type 3: CC1 is highly extended and the angle made by ICD, the hinge and N-GFP is obtuse, with two N-GFPs contact each other; Type 4: the same as type 3 except two N-GFPs are separated (see also, Figure 7C, middle and right); Type 5: CC1 is stretched and almost straight, separating ICD and N-GFP more than 50 nm. Bar represents 20 nm.

**Figure 7–S2.**
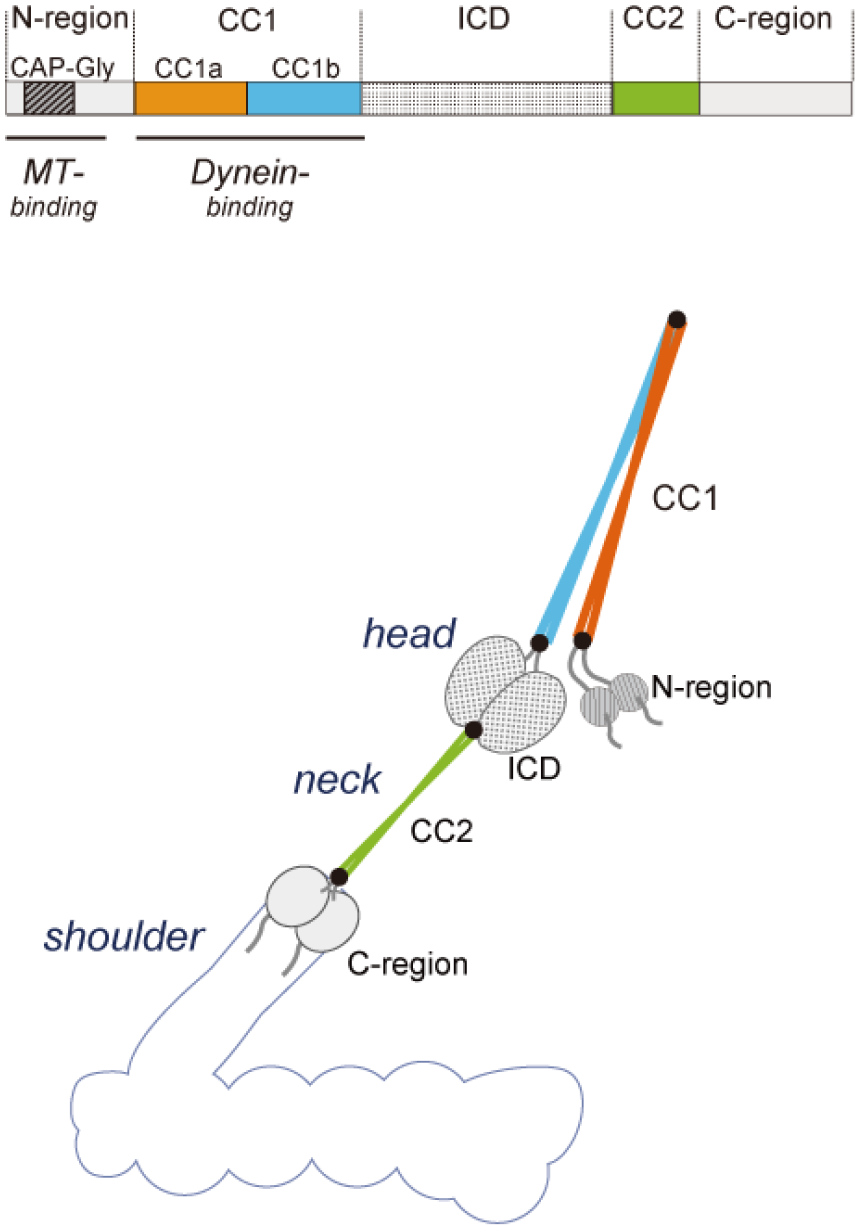
Domain organization of p150 A model for domain organization of p150. It illustrates the correspondence between five regions in the first amino acid sequence of p150 (N-region, CC1, ICD, CC2, C-region) and morphological domains observed by negative stain EM.

**Figure 7–S3.**
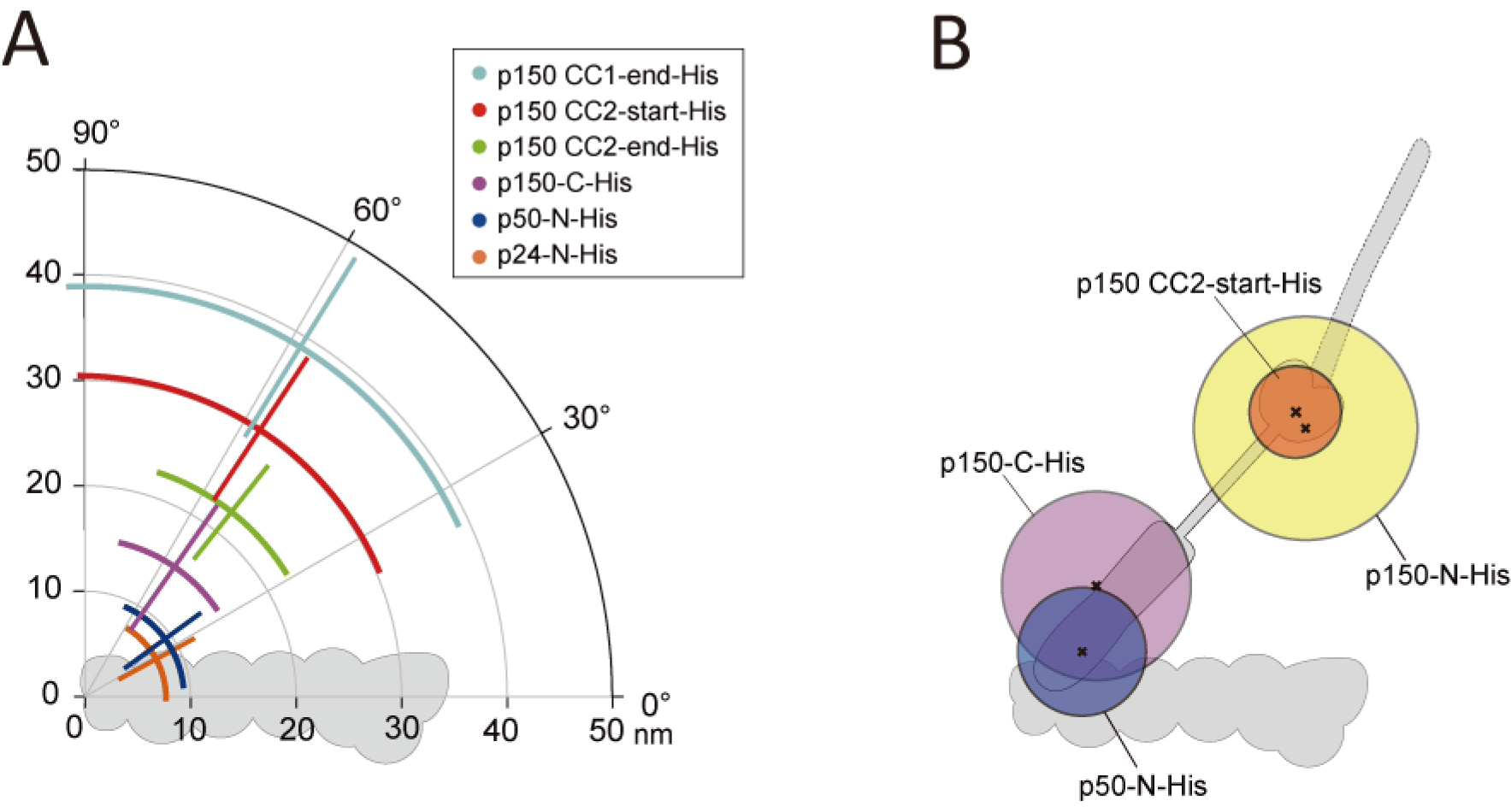
Distribution maps of the nanogold-labeled sites in the sidearm. (A) Summary of the nanogold labeling of representative mutants. Here, the positions of the gold nanoparticles were measured from the barbed-end as in Figure 1–S3A, right, blue graph. Note that, for the p150 CC1-end-His and the p150 CC2-start-His, the origins were set at the barbed-end of the Arp1 rod, which was different from the previous measurements (Figure 3D and Figure 4B, respectively). The radial and angular lines represent the SD of the measurement and their cross points indicate the mean value (listed in Table S2). (B) The range of distributions of gold nanoparticles for four representative mutants. SDs of the nanogold distributions for four mutants, calculated by root mean squared distance, are shown as radii of circles around the centroids (listed in Table S2). Original plot data were shown in Figures 2C, 2E, 4B and 5B for p150-N-His, p150-C-His, p150 CC2-start-His and p50-N-His, respectively.

**Figure 7–S4.**
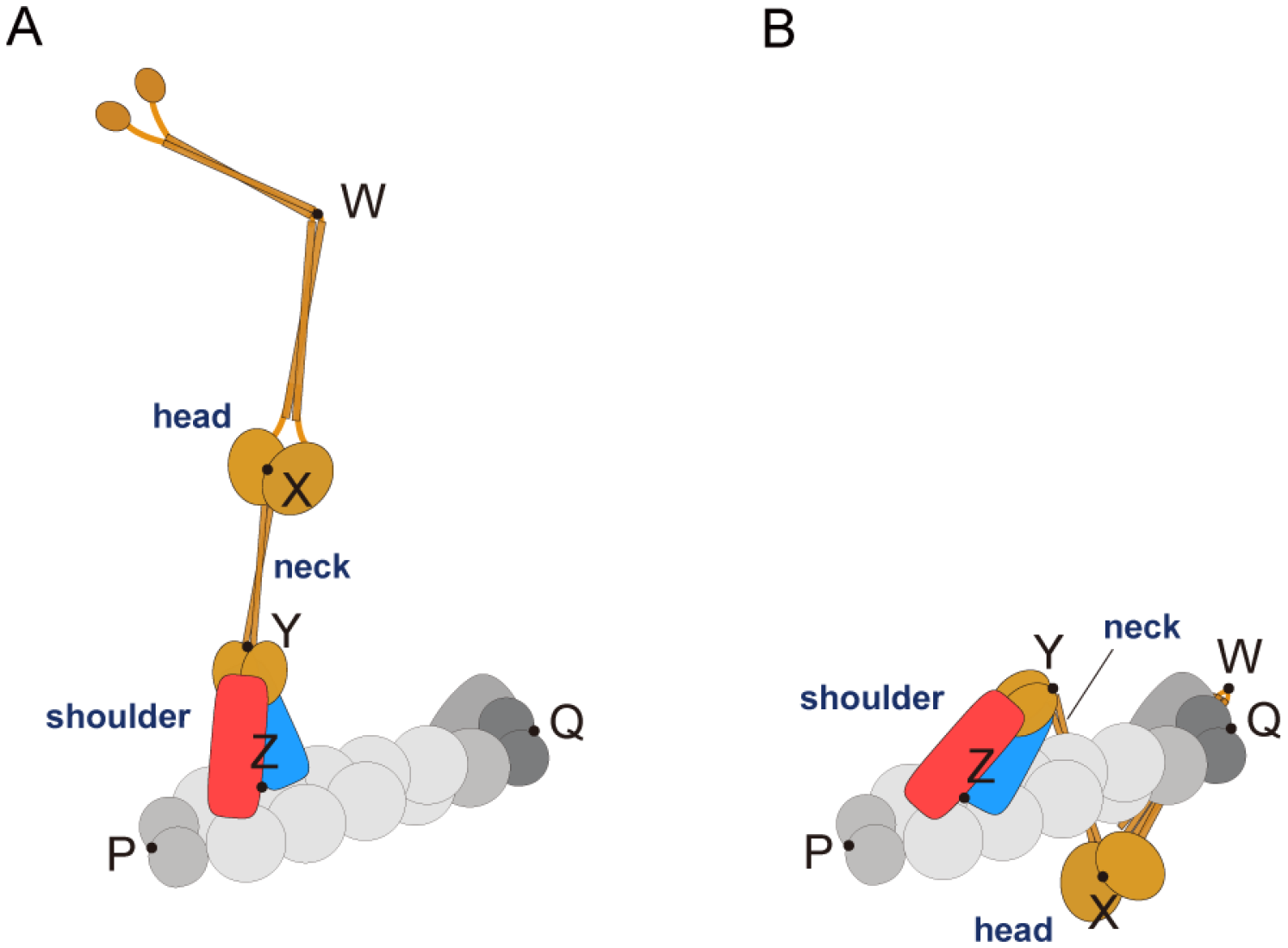
Models of undocked and docked sidearm. (A) A model of dynactin with undocked sidearm. CC1 is depicted as in the extended form. The points P, Q, W—Z indicate the reference points defined in Figure 1–S3. When sidearm is undocked, the p50-p24 arms in the shoulder domain (blue and red, corresponding to Arm-1 and Arm-2 in Urnavicius et al. (2015), respectively) are supposed to move flexibly (Figure 1B, **green**) as seen in the molecules (a)—(h) in Figure 1. (B) A model of dynactin with docked sidearm. CC1 is depicted as in the folded form and as it docks to the pointed end of the Arp1 rod, reflecting the structure of Urnavicius et al. (2015). When sidearm is docked, the p50-p24 arms are supposed to be confined along the Arp1 rod (Figure 1B, **red**) as seen in the molecule (i) in Figure 1. Compare with the averaged structure of the shoulder domain in the previous cryo-EM images presumably in the docked form (Chowdhury et al., 2015; Urnavicius et al., 2015).

**Table S1.**
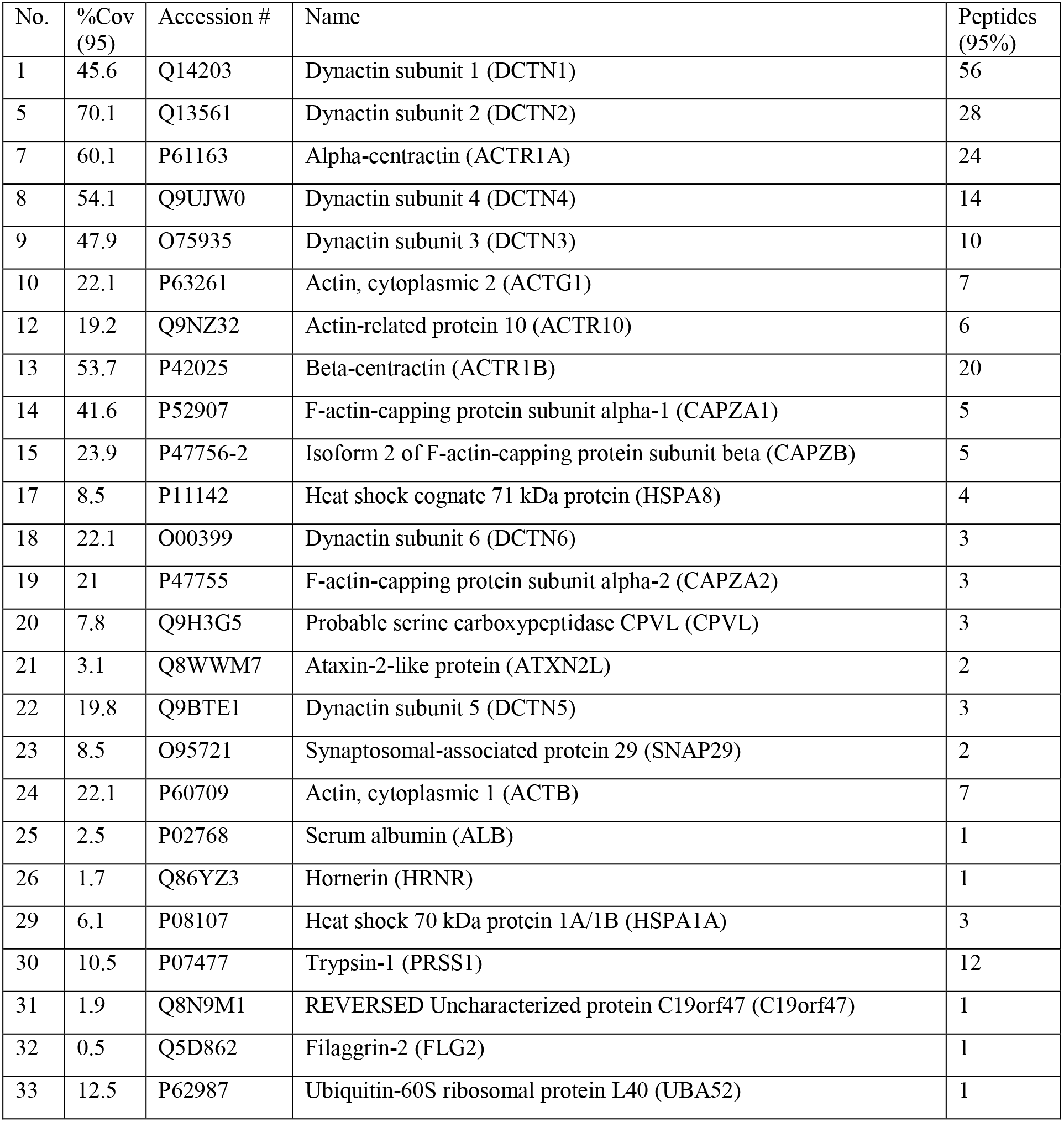
List of the proteins in the purified fraction of recombinant dynactin identified by LC’MS/MS spectrometry. Summary of proteins identified in the purified dynactin (SBP-p62). Keratin was removed from the list for simplification. %Cov (95) represents the percentage of matching amino acids from identified peptides having confidence greater than or equal to 95%, divided by the total number of amino acids in the sequence. Peptides (95%) represents the number of distinct peptide having 95% confidence.

| No. | %Cov (95) | Accession # | Name | Peptides (95%) |
| --- | --- | --- | --- | --- |
| 1 | 45.6 | Q14203 | Dynactin subunit 1 (DCTN1) | 56 |
| 5 | 70.1 | Q13561 | Dynactin subunit 2 (DCTN2) | 28 |
| 7 | 60.1 | P61163 | Alpha-centractin (ACTR1A) | 24 |
| 8 | 54.1 | Q9UJW0 | Dynactin subunit 4 (DCTN4) | 14 |
| 9 | 47.9 | O75935 | Dynactin subunit 3 (DCTN3) | 10 |
| 10 | 22.1 | P63261 | Actin, cytoplasmic 2 (ACTG1) | 7 |
| 12 | 19.2 | Q9NZ32 | Actin-related protein 10 (ACTR10) | 6 |
| 13 | 53.7 | P42025 | Beta-centractin (ACTR1B) | 20 |
| 14 | 41.6 | P52907 | F-actin-capping protein subunit alpha-1 (CAPZA1) | 5 |
| 15 | 23.9 | P47756-2 | Isoform 2 of F-actin-capping protein subunit beta (CAPZB) | 5 |
| 17 | 8.5 | P11142 | Heat shock cognate 71 kDa protein (HSPA8) | 4 |
| 18 | 22.1 | O00399 | Dynactin subunit 6 (DCTN6) | 3 |
| 19 | 21 | P47755 | F-actin-capping protein subunit alpha-2 (CAPZA2) | 3 |
| 20 | 7.8 | Q9H3G5 | Probable serine carboxypeptidase CPVL (CPVL) | 3 |
| 21 | 3.1 | Q8WWM7 | Ataxin-2-like protein (ATXN2L) | 2 |
| 22 | 19.8 | Q9BTE1 | Dynactin subunit 5 (DCTN5) | 3 |
| 23 | 8.5 | O95721 | Synaptosomal-associated protein 29 (SNAP29) | 2 |
| 24 | 22.1 | P60709 | Actin, cytoplasmic 1 (ACTB) | 7 |
| 25 | 2.5 | P02768 | Serum albumin (ALB) | 1 |
| 26 | 1.7 | Q86YZ3 | Hornerin (HRNR) | 1 |
| 29 | 6.1 | P08107 | Heat shock 70 kDa protein 1A/1B (HSPA1A) | 3 |
| 30 | 10.5 | P07477 | Trypsin-1 (PRSS1) | 12 |
| 31 | 1.9 | Q8N9M1 | REVERSED Uncharacterized protein C19orf47 (C19orf47) | 1 |
| 32 | 0.5 | Q5D862 | Filaggrin-2 (FLG2) | 1 |
| 33 | 12.5 | P62987 | Ubiquitin-60S ribosomal protein L40 (UBA52) | 1 |

**Table S2.** Positions of the labeled gold nanoparticles. Statistics on the centroid (Table A) and radius and angular (Table B) of the labeled gold. The positions of the gold nanoparticles were measured setting the origin at either the center of the head (head; red graph in Figure 1—S3A) or the barbed-end of the Arp1 rod (B-end; blue graph in Figure 1—S3A). SDs of the centroids in Table A were calculated by root mean squared distance to characterize dispersion around the centroid. Note that the position of centroid in Table A and mean values of the radius and angular in Table B were not identical in polar cordinates. For some mutants, both types of analyses were performed from the same nanogold labeling experimet but the final data set and number of particles analyzed could be different when origins were set differently (p150 CC1-end-His, p150 CC2-start-His and p150 CC2-end-His). For p150-N-His, p150 CC1-start-His, and p150 CC1-hinge-His, gold nanoparticles which did not make direct contact with the head domain were difficult to judge whether they were specifically bound to His-tag or not. We set the cut-off distance and only the gold nanoparticles within 20 nm from the center of the head were analyzed (p150-N-His and CC1-start-His) or only the molecules with the decernible protrusion (formed by CC1) were analyzed (p150-hinge-His).

| Table A | Constructs | r (nm) | $\theta$ (°) | SD (nm) | N | Origin | Fig. |
| --- | --- | --- | --- | --- | --- | --- | --- |
|  | p150-N-His | 2.8 | 51 | 10.7 | 74 | head | 2C,7S3B |
|  | p150-C-His | 14.2 | 54 | 9.0 | 98 | B-end | 2E,7S3B |
|  | p150 CC1-start-His | 2.2 | 12 | 10.6 | 76 | head | 3B, 3S1 |
|  | p150 CC1-hinge-His | 6.8 | 99 | 21.9 | 41 | head | 3B, 3S1 |
|  | p150 CC1-end-His | 4.6 | 179 | 7.2 | 110 | head | 3B, 3S1 |
|  | p150 CC2-start-His | 1.2 | 15 | 4.4 | 95 | head | 4B,7S3B |
|  | p150 CC2-end-His | 20.3 | 46 | 8.8 | 73 | B-end | 4D |
|  | p50-N-His | 8.3 | 36 | 6.1 | 64 | B-end | 5B,7S3B |
|  | p24-N-His | 6.9 | 37 | 5.5 | 32 | B-end | 5B |
| Table B | Constructs | r (nm) | $\theta$ (°) | – | N | Origin | Fig. |
|  | p150-C-His | 15.2 ± 7.3 | 55 ± 22 | – | 98 | B-end | 7S3A |
|  | p150 CC1-end-His | 39.2 ± 9.9 | 58 ± 34 | – | 94 | B-end | 7S3A |
|  | p150 CC2-start-His | 30.6 ± 8.1 | 57 ± 33 | – | 108 | B-end | 7S3A |
|  | p150 CC2-end-His | 22.5 ± 5.7 | 52 ± 20 | – | 95 | B-end | 7S3A |
|  | p50-N-His | 9.3 ± 4.5 | 36 ± 30 | – | 64 | B-end | 7S3A |
|  | p24-N-His | 7.8 ± 4.1 | 29 ± 31 | – | 32 | B-end | 7S3A |

mean ± SD

**Table S3.**
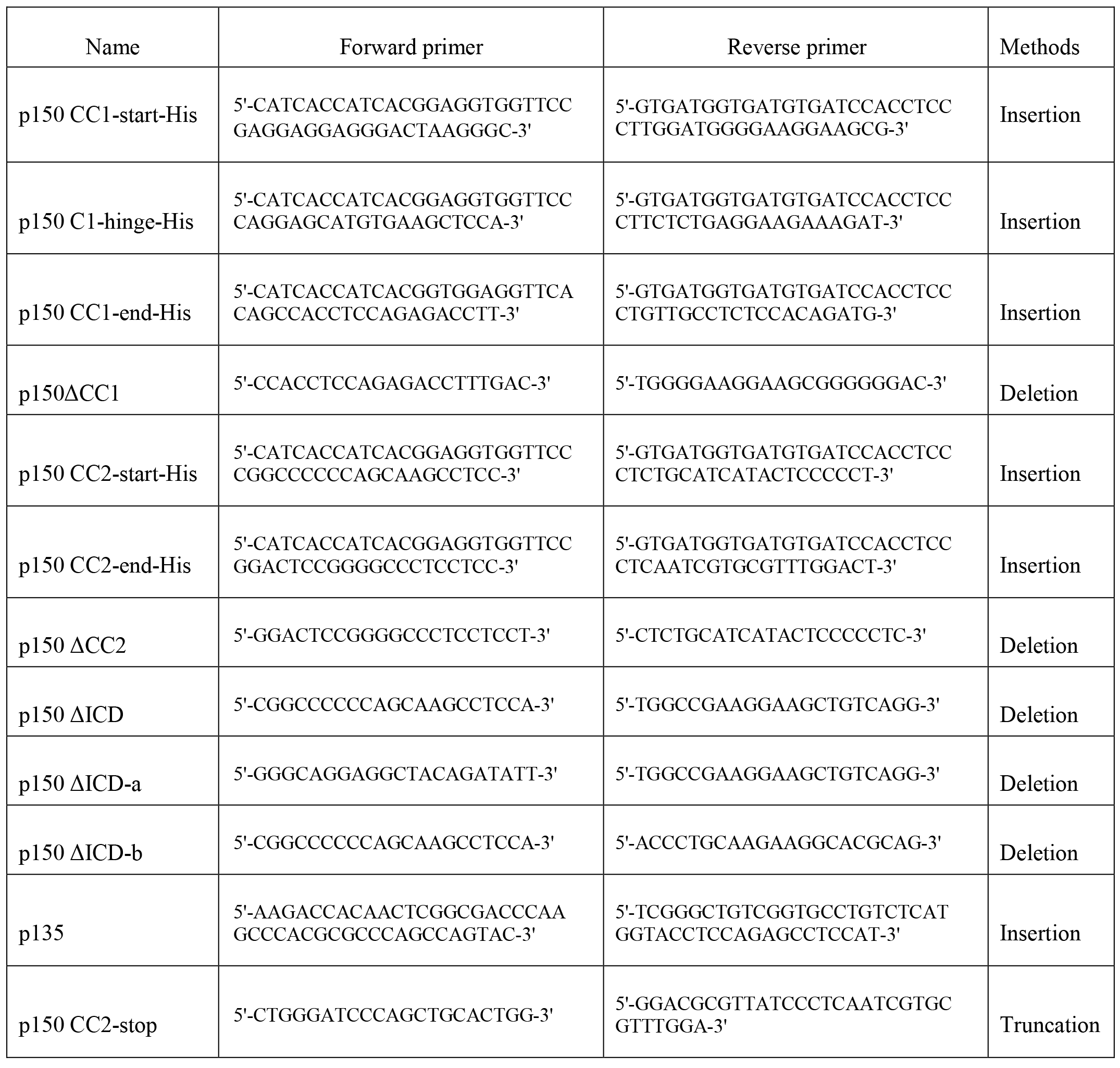
List of primers used for construction of p150 mutants.

## References

Akhmanova A, Steinmetz MO (2008) Tracking the ends: a dynamic protein network controls the fate of microtubule tips. Nat Rev Mol Cell Biol 9(4):309–22.

Ayloo S, Lazarus JE, Dodda A, Tokito M, Ostap EM, Holzbaur EL. (2014) Dynactin functions as both a dynamic tether and brake during dynein-driven motility. Nat Commun 5:4807.

Baumbach J, Murthy A, McClintock MA, Dix CI, Zalyte R, Hoang HT, Bullock SL (2017) Lissencephaly-1 is a context-dependent regulator of the human dynein complex. Elife 6: e21768.

Carter AP, Diamant AG, Urnavicius L (2016) How dynein and dynactin transport cargos: a structural perspective. Curr Opin Struct Biol 37:62–70.

Chowdhury S, Ketcham SA, Schroer TA, Lander GC (2015) Structural organization of the dynein-dynactin complex bound to microtubules. Nat Struct Mol Biol 22(4):345–7.

Cianfrocco MA, DeSantis ME, Leschziner AE, Reck-Peterson SL (2015) Mechanism and regulation of cytoplasmic dynein. Annu Rev Cell Dev Biol 31:83–108.

Culver-Hanlon TL, Lex SA, Stephens AD, Quintyne NJ, King SJ (2006) A microtubule-binding domain in dynactin increases dynein processivity by skating along microtubules. Nat Cell Biol 8(3): 264–270

Dixit R, Levy JR, Tokito M, Ligon LA, Holzbaur EL (2008) Regulation of dynactin through the differential expression of p150Glued isoforms. J Biol Chem 283(48): 33611–33619.

Echeverri CJ, Paschal BM, Vaughan KT, Vallee RB (1996) Molecular characterization of the 50-kD subunit of dynactin reveals function for the complex in chromosome alignment and spindle organization during mitosis. J Cell Biol 132(4): 617–633.

Eckley DM, Gill SR, Melkonian KA, Bingham JB, Goodson HV, Heuser JE, Schroer TA (1999) Analysis of dynactin subcomplexes reveals a novel actin-related protein associated with the arp1 minifilament pointed end. J Cell Biol 147(2): 307–320.

Fan SS, Ready DF (1997) Glued participates in distinct microtubule-based activities in Drosophila eye development. Development 124(8):1497–507.

Farrer MJ, Hulihan MM, Kachergus JM, Dachsel JC, Stoessl AJ, Grantier LL, Calne S, Calne DB, Lechevalier B, Chapon F, Tsuboi Y, Yamada T, Gutmann L, Elibol B, Bhatia KP, Wider C, Vilarino-Guell C, Ross OA, Brown LA, Castanedes-Casey M, Dickson DW, Wszolek ZK (2009) DCTN1 mutations in Perry syndrome. Nat Genet 41(2): 163–165.

Fujimura T, Shinohara Y, Tissot B, Pang PC, Kurogochi M, Saito S, Arai Y, Sadilek M, Murayama K, Dell A, Nishimura S, Hakomori SI (2008) Glycosylation status of haptoglobin in sera of patients with prostate cancer vs. benign prostate disease or normal subjects. Int J Cancer 122: 39–49.

Guesdon A, Bazile F, Buey RM, Mohan R, Monier S, García RR, Angevin M, Heichette C, Wieneke R, Tampé R, Duchesne L, Akhmanova A, Steinmetz MO, Chrétien D (2016) EB1 interacts with outwardly curved and straight regions of the microtubule lattice. Nat Cell Biol 18(10):1102–8.

Hayashi T, Saito T, Fujimura T, Hara K, Takamochi K, Mitani K, Mineki R, Kazuno S, Oh S, Ueno T, Suzuki K, Yao T. (2013) Galectin-4, a novel predictor for lymph node metastasis in lung adenocarcinoma. PLoS One 8(12):e81883.

Hodgkinson JL, Peters C, Kuznetsov SA, Steffen W (2005) Three-dimensional reconstruction of the dynactin complex by single-particle image analysis. Proc Natl Acad Sci U S A 102(10): 3667–3672.

Ichikawa M, Saito K, Yanagisawa HA, Yagi T, Kamiya R, Yamaguchi S, Yajima J, Kushida Y, Nakano K, Numata O, Toyoshima YY (2015) Axonemal dynein light chain-1 locates at the microtubule-binding domain of the y heavy chain. Mol Biol Cell 26(23):4236–47.

Ichikawa M, Watanabe Y, Murayama T, Toyoshima YY (2011) Recombinant human cytoplasmic dynein heavy chain 1 and 2: observation of dynein-2 motor activity in vitro. FEBS Lett 585(15): 2419–2423.

Imai H, Narita A, Schroer TA, Maeda Y (2006) Two-dimensional averaged images of the dynactin complex revealed by single particle analysis. J Mol Biol 359(4): 833–839.

Imai H, Narita A, Maéda Y, Schroer TA (2014) Dynactin 3D structure: implications for assembly and dynein binding. J Mol Biol 426(19): 3262–3271.

Ishihara K, Nguyen PA, Groen AC, Field CM, Mitchison TJ (2014) Microtubule nucleation remote from centrosomes may explain how asters span large cells. Proc Natl Acad Sci USA 111(50):17715–22.

Jacquot G, Maidou-Peindara P, Benichou S (2010) Molecular and functional basis for the scaffolding role of the p50/dynamitin subunit of the microtubule-associated dynactin complex. J Biol Chem 285(30): 23019–23031.

Jha R, Roostalu J, Cade NI, Trokter M, Surrey T (2017) Combinatorial regulation of the balance between dynein microtubule end accumulation and initiation of directed motility. EMBO J 36(22):3387–3404.

Kardon JR, Reck-Peterson SL, Vale RD (2009) Regulation of the processivity and intracellular localization of Saccharomyces cerevisiae dynein by dynactin. Proc Natl Acad Sci U S A 106(14): 5669–5674.

Kardon JR, Vale RD (2009) Regulators of the cytoplasmic dynein motor. Nat Rev Mol Cell Biol 10(12): 854–865.

Karki S, Holzbaur EL (1995) Affinity chromatography demonstrates a direct binding between cytoplasmic dynein and the dynactin complex. J Biol Chem 270(48): 28806–28811.

King SJ, Brown CL, Maier KC, Quintyne NJ, Schroer TA (2003) Analysis of the dynein-dynactin interaction in vitro and in vivo. Mol Biol Cell 14(12): 5089–5097.

Kitai T, Watanabe Y, Toyoshima YY, Kobayashi T, Murayama T, Sakaue H, Suzuki H, Takahagi T (2011) Simple Method of Synthesizing Nickel-Nitrilotriacetic Acid Gold Nanoparticles with a Narrow Size Distribution for Protein Labeling. Japanese Journal of Applied Physics 50: 095002–095005. http://dx.doi.org/10.1143/JJAP.50.095002

Kobayashi T, Shiroguchi K, Edamatsu M, Toyoshima YY (2006) Microtubule-binding properties of dynactin p150 expedient for dynein motility. Biochem Biophys Res Commun 340(1): 23–28.

Kobayashi T, Morone N, Kashiyama T, Oyamada H, Kurebayashi N, Murayama T (2008) Engineering a novel multifunctional green fluorescent protein tag for a wide variety of protein research. PLoS One 3(12):e3822.

Kobayashi T, Murayama T (2009) Cell cycle-dependent microtubule-based dynamic transport of cytoplasmic dynein in mammalian cells. PLoS One 4(11): e7827.

Kobayashi T, Miyashita T, Murayama T, Toyoshima YY (2017) Dynactin has two antagonistic regulatory domains and exerts opposing effects on dynein motility. PLoS One 12(8):e0183672.

Lloyd TE, Machamer J, O’Hara K, Kim JH, Collins SE, Wong MY, Sahin B, Imlach W, Yang Y, Levitan ES, McCabe BD, Kolodkin AL (2012) The p150(Glued) CAP-Gly domain regulates initiation of retrograde transport at synaptic termini. Neuron 74(2):344–60.

Maier KC, Godfrey JE, Echeverri CJ, Cheong FK, Schroer TA (2008) Dynamitin mutagenesis reveals protein-protein interactions important for dynactin structure. Traffic 9(4): 481–491.

Mcgrail M, Gepner J, Silvanovich A, Ludmann S, Serr M, Hays TS (1995) Regulation of Cytoplasmic Dynein Function In Vivo by the Drosophila Glued complex. J. Cell Biol. 131(2): 411–425.

McGuffin LJ, Bryson K, Jones DT (2000) The PSIPRED protein structure prediction server. Bioinformatics 16(4): 404–405.

McKenney RJ, Huynh W, Vale RD, Sirajuddin M (2016) Tyrosination of a-tubulin controls the initiation of processive dynein-dynactin motility. EMBO J 35(11):1175–85.

Melkonian KA, Maier KC, Godfrey JE, Rodgers M, Schroer TA (2007) Mechanism of dynamitin-mediated disruption of dynactin. J Biol Chem 282(27): 19355–19364.

Miura M, Matsubara A, Kobayashi T, Edamatsu M, Toyoshima YY (2010) Nucleotide-dependent behavior of single molecules of cytoplasmic dynein on microtubules in vitro. FEBS Lett 584(11): 2351–2355.

Morgan JL, Song Y, Barbar E (2011) Structural dynamics and multiregion interactions in dynein-dynactin recognition. J Biol Chem 286(45): 39349–39359.

Moughamian AJ, Holzbaur EL (2012) Dynactin is required for transport initiation from the distal axon. Neuron 74(2):331–43.

Puls I, Jonnakuty C, LaMonte BH, Holzbaur EL, Tokito M, Mann E, Floeter MK, Bidus K, Drayna D, Oh SJ, Brown RH, Jr., Ludlow CL, Fischbeck KH (2003) Mutant dynactin in motor neuron disease Nat Genet 33(4): 455–456.

Quintyne NJ, Gill SR, Eckley DM, Crego CL, Compton DA, Schroer TA (1999) Dynactin is required for microtubule anchoring at centrosomes. J Cell Biol 147(2):321–34.

Schafer DA, Gill SR, Cooper JA, Heuser JE, Schroer TA (1994) Ultrastructural analysis of the dynactin complex: an actin-related protein is a component of a filament that resembles F-actin. J Cell Biol 126(2): 403–412.

Schroer TA (2004) Dynactin. Annu Rev Cell Dev Biol 20: 759–779.

Siglin AE, Sun S, Moore JK, Tan S, Poenie M, Lear JD, Polenova T, Cooper JA, Williams JC (2013) Dynein and dynactin leverage their bivalent character to form a high-affinity interaction. PLoS One 8(4): e59453.

Song K, Awata J, Tritschler D, Bower R, Witman GB, Porter ME, Nicastro D (2015) In situ localization of N and C termini of subunits of the flagellar nexin-dynein regulatory complex (N-DRC) using SNAP tag and cryo-electron tomography. J Biol Chem 290(9):5341–53.

Steinmetz MO, Akhmanova A (2008) Capturing protein tails by CAP-Gly domains. Trends Biochem Sci 33(11): 535–545.

Suzuki K, Miyazaki M, Takagi J, Itabashi T, Ishiwata S (2017) Spatial confinement of active microtubule networks induces large-scale rotational cytoplasmic flow. Proc Natl Acad Sci USA 114(11):2922–2927.

Swaroop A, Paco-Larson ML, Garen A (1985) Molecular genetics of a transposon-induced dominant mutation in the Drosophila locus Glued. Proc Natl Acad Sci USA 82(6):1751–5.

Tokito MK, Howland DS, Lee VM, Holzbaur EL (1996) Functionally distinct isoforms of dynactin are expressed in human neurons. MolBiol Cell 7(8): 1167–1180.

Tripathy SK, Weil SJ, Chen C, Anand P, Vallee RB, Gross SP (2014) Autoregulatory mechanism for dynactin control of processive and diffusive dynein transport. Nat Cell Biol. 16(12):1192–1201.

Urnavicius L, Zhang K, Diamant AG, Motz C, Schlager MA, Yu M, Patel NA, Robinson CV, Carter AP. (2015) The structure of the dynactin complex and its interaction with dynein. Science 347(6229):1441–1446.

Vaughan KT, Vallee RB (1995) Cytoplasmic dynein binds dynactin through a direct interaction between the intermediate chains and p150Glued. J Cell Biol 131(6 Pt 1): 1507–1516.

Waterman-Storer CM, Karki S, Holzbaur EL (1995) The p150Glued component of the dynactin complex binds to both microtubules and the actin-related protein centractin (Arp-1). Proc Natl Acad Sci U S A 92(5): 1634–1638.

Weisbrich A, Honnappa S, Jaussi R, Okhrimenko O, Frey D, Jelesarov I, Akhmanova A, Steinmetz MO (2007) Structure-function relationship of CAP-Gly domains. Nat Struct Mol Biol 14(10): 959–967.

Yeh TY, Quintyne NJ, Scipioni BR, Eckley DM, Schroer TA (2012) Dynactin’s pointed-end complex is a cargo-targeting module. Mol Biol Cell 23(19): 3827–3837.

Yeh TY, Kowalska AK, Scipioni BR, Cheong FK, Zheng M, Derewenda U, Derewenda ZS, Schroer TA (2013) Dynactin helps target Polo-like kinase 1 to kinetochores via its left-handed beta-helical p27 subunit. EMBO J 32(7): 1023–1035.

Zhapparova ON, Bryantseva SA, Dergunova LV, Raevskaya NM, Burakov AV, Bantysh OB, Shanina NA, Nadezhdina ES (2009) Dynactin subunit p150Glued isoforms notable for differential interaction with microtubules. Traffic 10(11): 1635–1646.

